# Control adjustment costs limit goal flexibility: Empirical evidence and a computational account

**DOI:** 10.1101/2023.08.22.554296

**Authors:** Ivan Grahek, Xiamin Leng, Sebastian Musslick, Amitai Shenhav

**Author notes:** These authors contributed equally to this work (Ivan Grahek, Xiamin Leng). This work was supported by the Training Program for Interactionist Cognitive Neuroscience T32-MH115895 (X.L.), NIMH R01MH124849 (A.S.), NIMH R21MH122863 (A.S.), NSF Career Award 204611 (A.S.), and NIH Award 1S10OD02518. The authors thank Jonathan D. Cohen, Alexander Fengler, Atsushi Kikumoto, Harrison Ritz, Ainsley Bonin, and all members of the Shenhav lab for many fruitful discussions of this work. We thank Mahalia Prater Fahey and Debbie Yee for providing the datasets on which some of the ideas in this manuscript were initially tested. Raw data and analysis scripts are available on GitHub (https://github.com/igrahek/ControlAdjustmentCosts.git). This study was not preregistered. Corresponding authors. E-mail address &.

## Abstract

A cornerstone of human intelligence is the ability to flexibly adjust our cognition and behavior as our goals change. For instance, achieving some goals requires efficiency, while others require caution. Different goals require us to engage different control processes, such as adjusting how attentive and cautious we are. Here, we show that performance incurs control adjustment costs when people adjust control to meet changing goals. Across four experiments, we provide evidence of these costs, and validate a dynamical systems model explaining the source of these costs. Participants performed a single cognitively demanding task under varying performance goals (e.g., being fast or accurate). We modeled control allocation to include a dynamic process of adjusting from one’s current control state to a target state for a given performance goal. By incorporating inertia into this adjustment process, our model accounts for our empirical finding that people under-shoot their target control state more (i.e., exhibit larger adjustment costs) when goals switch rather than remain fixed (Study 1). Further validating our model, we show that the magnitude of this cost is increased when: distances between target states are larger (Study 2), there is less time to adjust to the new goal (Study 3), and goal switches are more frequent (Study 4). Our findings characterize the costs of adjusting control to meet changing goals, and show that these costs emerge directly from cognitive control dynamics. In so doing, they shed new light on the sources of and constraints on flexibility of goal-directed behavior.

## Introduction

Every day we pursue hundreds of goals. As we make coffee, answer emails, and attend meetings, we continuously adjust how we process information and act. Achieving these varied goals requires profound changes in how we engage with the tasks before us. For example, some goals are achieved with minimal focus (e.g., e-mailing a friend), while others require greater caution (e.g., e-mailing our boss). Goal changes thus require adjustments of cognitive control (e.g., levels of attention and caution), which are known to come at a cost: we are better at achieving our goals in isolation than in situations that demand frequent goal changes. Thus, control adjustment costs limit our ability to flexibly move between goals, but the source of these costs remains unknown.

Cognitive control processes allow us to align information processing with our current goal, enabling us to achieve our goals by focusing on goal-relevant information and suppressing distractions (Miyake et al., 2000; Diamond, 2013; Botvinick & Cohen, 2014). Critically, changes in our environment (e.g., increased incentives) or task instructions (e.g., needing to complete a task quickly) induce adjustments of continuous cognitive control states that determine how we engage with our current task. For example, people allocate more attention when expected incentives or task demands are high (Botvinick & Braver, 2015; Shenhav et al., 2017; Parro et al., 2018). They also increase levels of attention and caution after they commit errors or experience response conflict (Egner, 2007; Danielmeier & Ullsperger, 2011), and they adjust control when task instructions change (Ratcliff & Rouder, 1998; Forstmann et al., 2010). Such control adjustments are ubiquitous and have inspired theoretical models aimed at understanding how people determine which levels of control to engage (Botvinick et al., 2001; Bogacz et al., 2006; Ritz et al., 2022). However, despite the core role of control adjustments in goal flexibility, the dynamics of transitions between different continuous control states, and the potential costs of such transitions, remain unexplored.

Relevant to these open questions, a substantial body of work has examined the costs of control adjustment under the rubric of task-switching costs. In this research, participants transition between different task sets (e.g., classifying a digit as odd/even or greater/less than 5), requiring them to change which information they attend to (e.g., which stimulus feature) and/or the corresponding response mapping (Allport et al., 1994; Rogers & Monsell, 1995; MacDowell et al., 2022; Frings et al., 2020). This work has consistently demonstrated that participants are impaired at achieving a given task goal on trials immediately following a task switch, relative to those involving a repeat of the previous task (switch costs; Kiesel et al., 2010; Koch et al., 2018). While this research has provided critical insight into constraints on control flexibility, it has been primarily limited to understanding changes in control when transitioning between discrete task sets, which can entail changes in levels of control, but also changes in the information being processed and corresponding response contingencies. Moreover, as participants transition between these tasks their performance goal typically remains the same (e.g., to be fast and accurate). To better understand the costs of adjusting control states along a continuum, we sought to model and test when and how such costs arise when goals change in the absence of task changes.

Here, we develop a dynamical systems model that characterizes environments in which a person transitions between different performance goals (e.g., performing a task quickly or accurately), and must adjust their control state to best meet their current goal. This model predicts that people will settle on a “target” control state (e.g., high level of focus and caution) when continually pursuing a given performance goal (e.g., “be accurate”), and will settle on a different target control state (e.g., low level of focus and caution) when continually pursuing a different performance goal (e.g., “be fast”). Critically, the model includes an inertia in transitions between control states. Thus, the model predicts that when transitioning between performance goals people will undershoot these target states. We will refer to this undershoot as a *control adjustment cost*. This cost arises from control dynamics in the sense that control states are not immediately adjusted following a goal change, but that adjustments are gradual and unfold over time. Thus, the model predicts the existence of periods in which there is a mismatch between the desired and the actual control state. This is reflected in the adjustment cost, and the model predicts that the cost emerges are a function of: (1) the need to adjust control states, (2) the distance that needs to be traversed in control space, (3) the time allowed for transitions, and (4) the frequency of goal switches. To test these predictions, we have participants across four experiments perform a cognitive task in which stimulus-response contingencies are held constant, but the instructed performance goal varies (e.g., either to perform the task quickly or accurately). Our findings confirm all four predictions of our model, and collectively suggest that the costs associated with switching between goals emerge from the dynamics of cognitive control adjustments which are gradual and characterized by inertia.

## Results

### Control Adjustment Costs: A Dynamical Systems Model

Consider an environment in which participants perform a control-demanding task such as naming the ink color of a color word (Figure 1A). While they continue performing that task, their instruction for how to perform the task (their *performance goal*) varies: sometimes they are asked to perform the task as accurately as possible (Accuracy goal); other times they are asked to perform the as fast as possible (Speed goal). Adjusting to either goal will require participants to reconfigure their information processing accordingly. Such an adjustment can be operationalized as dynamic switch between control states (e.g., low level of caution in the speed goal vs. high level of caution in the accuracy goal). Building on previous work in switching between discrete control states (Ueltzhöffer et al., 2015; Musslick, Bizyaeva, et al., 2019; Steyvers et al., 2019), we formalize this process as a dynamical system in which control is gradually adjusted from one control state (e.g., low decision threshold) to another (e.g., high decision threshold). This model’s key assumption derives from the concept of task-set inertia, which has been used to explain performance costs associated with goal switches between task sets (Allport et al., 1994; Gilbert & Shallice, 2002), but has never been evaluated in the context of switching between continuous control states such as levels of attention or response caution. The intuition is that such inertia in control adjustments will produce suboptimal behavior when people frequently switch between different control states. In order to simulate behavior arising from this model, and compare it to empirical data, we formalize the model as follows.

**Figure 1.**
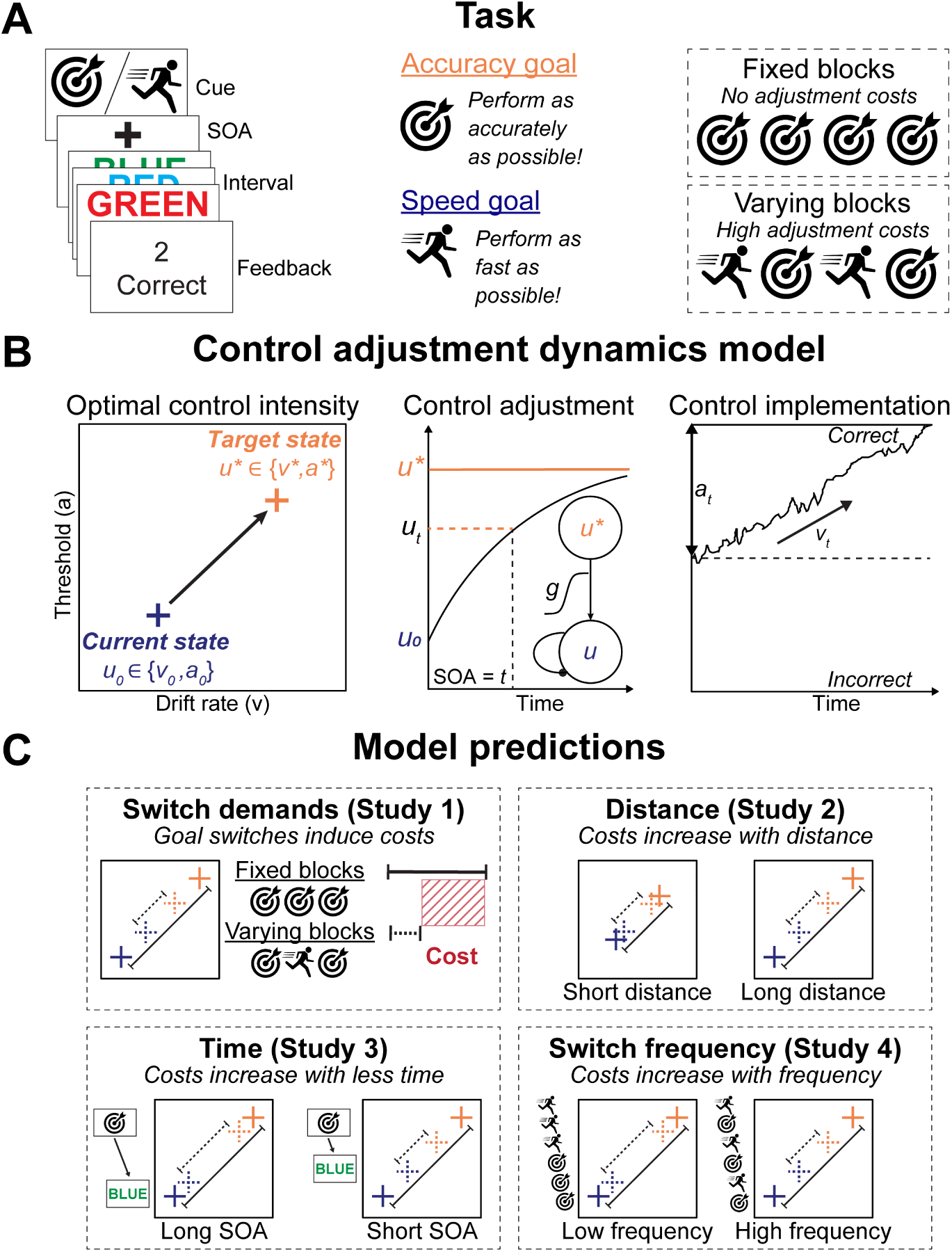
Behavioral task and the model of control dynamics. **A.** Participants performed an interval-based version of the Stroop task (left panel) in which they had a fixed amount of time to complete as many trials as they wished. Prior to each interval, they saw a cue (middle panel) instructing them about the current performance goal (e.g., speed or accuracy). Each experiment consisted of blocks (right panel) in which the performance goal never changed (fixed) and blocks in which the performance goals frequently changed (varying). **B.** The model assumes that once the performance goal changes (e.g., from Speed to Accuracy), control state (drift rate & threshold) must be adjusted from the current control state (blue) to the target state required by the new goal (orange). This adjustment happens gradually, starting at the onset of the goal cue, and stops once the model is presented with the first Stroop stimulus. Critically, the optimal control state might not be reached during within this time (SOA in the middle panel). Control adjustment dynamics depend on the distance between the current and the target state, which can also be expressed as a leaky accumulation based on the target state input. Once the task starts, the model implements the current control configuration to determine response selection and reaction time. **C.** The model makes four core predictions, which we test across four studies. First, the model predicts that environments that demand goal switches (varying blocks) will incur greater control adjustment costs than those with consistent goals (fixed blocks) (Study 1). Due to gradual control adjustments, this cost is observed as reduced distance between control states in varying relative to fixed blocks. The model further predicts that the magnitude of control adjustment costs increases with greater distance between control states (Study 2), less time to adjust control (Study 3), and increased frequency of goal switches (Study 4).

First (Figure 1B-left), the target control state depends on the current goal which in turn depends on the current incentives or performance goals. That target control state could be identical to or differ from the previous control state (e.g., the settings used for the most recent trial). Second (Figure 1B-middle), to the extent these states differ, the previous control state is gradually adjusted toward the target state. Third (Figure 1B-right), when the task onsets, the control state at that moment is used to determine response selection; critically, this control state that is ultimately implemented on a given trial need not match either the previous control state or the target state. We next describe each of these processing stages and model components in turn.

### Determining Control Signals for Each Goal

Control states are not directly observable, and thus studying adjustments in control requires a process model that allows for the specification of control signals. In the case of task goals that pertain to speed and/or accuracy, control signals are commonly operationalized through sequential sampling models of decision-making (Ratcliff & Rouder, 1998; Bogacz et al., 2006; Ratcliff & McKoon, 2008; Ritz et al., 2022). The Drift Diffusion Model is a model of decision-making that casts decision-making as a noisy accumulation of evidence towards one of the response boundaries. The speed of evidence accumulation (*drift rate* parameter) relates to processing efficiency, which can be modulated by attention. Higher drift rates lead to faster and more accurate responses. Response caution (*threshold* parameter) represents how much evidence is needed to make a decision. Higher thresholds are related to slower, but more accurate responses. Following our previous work (Leng et al., 2021; Grahek et al., 2023), we interpret levels of drift rate and threshold as markers of cognitive control levels. As detailed in the results section, we also control for variability in drift rate as a function of response congruency, which has been viewed as an alternative metric of control levels, but which can also primarily reflect stimulus-driven processes rather than control per se (Musslick, Cohen, et al., 2019).

Levels of drift rate and threshold represent two dimensions within a multivariate space of possible control signals (Bogacz et al., 2006; Musslick et al., 2015; Ritz et al., 2022). Computational models of cognitive control can identify a point in this multidimensional space (a *control state*) that maximizes one’s current expected benefits while minimizing control costs. To identify optimal control states for the two performance goals described above (Accuracy vs. Speed Goal), we can leverage a normative model proposed by Bogacz and colleagues (2006), that was recently further developed and empirically tested by Leng and colleagues (2021). This model finds optimal control states by maximizing the expected reward rate for different combinations of drift rates and thresholds (see the Methods section for details). We use this model to specify optimal control states (*v*^∗^, *a*^∗^) given the current state of the environment (including current performance goals; Figure 1B-left). These optimal control states can be understood as representing the target state for a given performance goal (e.g., when given the Speed goal), but how it is that a person shifts between control states remains unspecified.

### Cognitive Control Dynamics

When the environment changes (e.g., a change in the explicit performance goal or in the incentives for the current goal), the current control state needs to be adjusted to a new target state in order to perform optimally on the task (Figure 1B-middle). We formalize this idea by proposing that a control state at a given time (*u*_*t*_) is gradually adjusted towards the new target state (*u*^∗^). Here we consider two control signals: drift rate (*v*_*t*_) and threshold (*a*_*t*_) which are adjusted towards the new target states described above (*u*^∗^ ∈ {*v*^∗^, *a*^∗^}. We represent these control states as active units in a dynamical system in which activity states correspond to the activity of control units in a neural system (cf. Ueltzhöffer et al., 2015; Musslick et al., 2018, 2019). We simulate two control units whose activity at time *t* corresponds to the current level of drift rate *v*_*t*_ and threshold *a*_*t*_. Building on previous work in task-switching (Musslick, Bizyaeva, et al., 2019; Steyvers et al., 2019), the dynamics of control adjustments in this system are governed by an inertia parameter (τ) and by the distance between the current goal state (*u*_*t*_) and the target goal state (*u*^∗^):

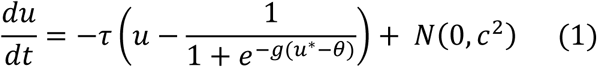

This model can be rewritten in the following way and understood as a leaky accumulation (Usher & McClelland, 2001) of the current control state:

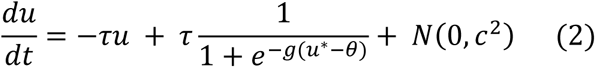

Target control state (*u*^∗^) represents the input to the system and is passed through a sigmoid activation function. The sigmoid activation function restricts the control intensity between the values of 0 and 1, and is offset by a bias term θ. The gain *g* regulates the sensitivity of the current processing intensity to changes in the input (e.g., changes in target states). This gain parameter has been suggested to modulate neural dynamics, similar to the effects of neuromodulators (Servan-Schreiber et al., 1990; Liljenström, 2003; Cools, 2015), in order to control the balance between stability and flexibility in the system (Musslick, Bizyaeva, et al., 2019; Musslick et al., 2018). The inertia parameter τ determines the rate of change in the input, which is determined by two factors: (1) a decay term that scales with current control level *u*_*t*_, and (2) an excitatory term that represents the optimal control level *u*^∗^ for that control unit. Consequently, the equilibrium state of the control unit’s net input is achieved when its control level matches the target control level (i.e., the sigmoidal transformation of *u*^∗^). Noise is zero-mean Gaussian with variance set by *c*. The dynamics dictated above highlight two key factors that will determine the degree to which one’s control state shifts between the current and the target state: distance between the two states (e.g., how different the current and target drift rates and thresholds are) and time to adjust (e.g., the time between being presented with the performance goal and task onset).

### Control Implementation

Once the adjustment of control signals is terminated (e.g., task onset), the current control state is implemented and used to generate behavior (Figure 1B-right). Based on this control state (*v*, *a*) and the features of the stimulus (e.g., whether the word and ink color indicate congruent or incongruent responses), a distribution of expected responses and response time likelihoods can be estimated from the Drift Diffusion Model (Navarro & Fuss, 2009).

### Model Predictions

The core assumption of the proposed model is that adjustments in control states (attention and caution) are subject to inertia (Figure 1C). Such inertia can prevent the control system from reaching a target control state. We refer to this undershoot in control adjustment as a *control adjustment cost*, and quantify this as the distance between the control state one achieves on a given trial and the inferred target control state for that trial (i.e., the control state one would settle on in the absence of a need for adjustment). This model makes a series of falsifiable predictions, which we systematically test in the studies reported below.

First, relative to task environments that maintain the same performance goal (and thus no need for control adjustment), environments that require regular changes in performance goals (and thus changes in control states) should result in smaller control adjustments on average (i.e., should incur larger adjustment costs) (Study 1). Second, participants should incur less of an adjustment cost when switching between two target states that are closer to one another in control space (e.g., based on the similarity of the corresponding performance goals) and more of an adjustment cost when those target states are further apart (Study 2). Third, participants should incur more of an adjustment cost the less time they are given to transition between their current control state and a different target state (e.g., based on the relative onset of goal and task stimulus information) (Study 3). Finally, participants should incur greater control adjustment costs the more frequently they are required to transition between control states (Study 4).

### Adjustment Costs Arise Under Conditions of Changing Performance Goals (Study 1)

Participants in Study 1 (N = 44) performed the color-word Stroop task during fixed time intervals (8-12s). On each interval, they were instructed to perform the task as fast (Speed goal) or as accurately (Accuracy goal) as possible. They did this in blocks in which the performance goal was either always the same (Fixed blocks), or the two goals alternated in random fashion (Varying blocks).

#### Model Predictions

Our model predicts that there is inertia in the adjustments of control signal, which will lead to different behavioral profiles in situations in which participants always have the same control signals compared to situations in which they have to frequently adjust control (Figure 1C). Thus, the model predicts that control intensities for the speed and accuracy conditions will differ between fixed and varying blocks, and that these will manifest in changes to drift rates and thresholds.

To formalize model predictions, we first simulated what level of drift rates and thresholds participants would set in the speed and accuracy conditions if there were no costs to switching between them (i.e., if one of them was their only goal; details in Methods section). We proceeded to simulate which levels of drift rate and threshold will be reached on average in blocks in which participants had to frequently switch between different goals (varying blocks). These simulations assumed that participants tried to reach the same levels of drift rates and thresholds as in fixed blocks, but failed to do so because they had to frequently adjust control states. Thus, control adjustment costs emerge, and they are revealed by the prediction that the distance between two control states is reduced in varying compared to fixed blocks (Figure 2B-left). This effect is generated by the need to frequently adjust control signals (move in the 2-D control space) in the varying blocks. Because of the gradual adjustment of control assumed by our model, such movement means that the control system will often be in between the two target states, rather than at the target state required by the current goal. Thus, the second prediction of our model (Figure 2D-left) is that in varying blocks control intensities for the two goals will be closer together on switch intervals (in which the agent has a different goal state compared to the previous interval) relative to the repeat intervals (in which the agent has the same goal state as on the previous interval).

**Figure 2.**
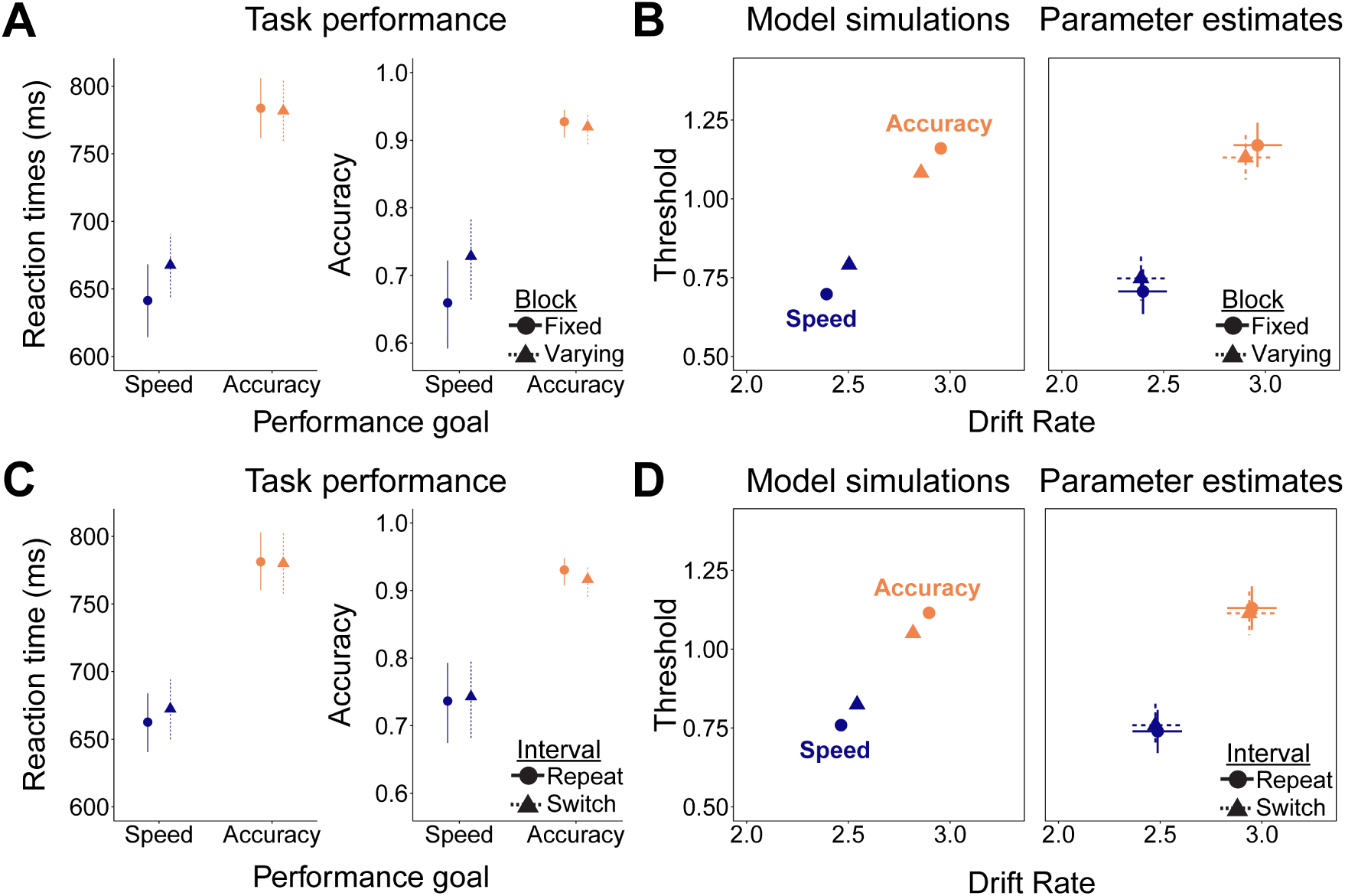
Frequent adjustments of control intensity impact behavioral performance and control intensity. **A.** In the speed condition (blue) participants were faster and less accurate in blocks with the fixed goal (solid) compared to blocks in which the goals were varying (dashed). Dots represent regression estimates and error bars 95% credible intervals. **B.** Model simulations (left) and drift diffusion parameter estimates (right) show that control intensities for the two performance goals are pulled closer in varying blocks which include goal switches. Error bars represent 95% credible intervals. **C.** In varying blocks, participants were faster in the speed condition when the previous interval had the speed (Repeat) relative to accuracy (Switch) goal. In the accuracy condition, they were less accurate in the switch relative to speed condition. **D.** Model simulations and parameter estimates for the switch and repeat intervals in varying blocks show that the estimates are pulled closer together on switch relative to repeat intervals.

We tested model predictions by first comparing the behavioral performance of our participants on fixed and varying blocks, as well as on switch and repeat intervals within the varying blocks. Then, we fit the Drift Diffusion Model to behavioral data in order to obtain the drift rates and thresholds across conditions and compare them to model predictions. Across these analyses, the key index of control adjustment costs is the extent to which people adjust their control state less between the two performance goals when they are in varying relative to fixed blocks. Control adjustment costs are evidenced by decreased difference between Speed and Accuracy conditions when participants frequently switch between these two performance goals. We tested for these costs based on differences in raw behavioral performance across the conditions (accuracies and RTs; *model-agnostic* indices of control adjustment costs) and based on control state estimates from fitting behavioral data to a Drift Diffusion Model (*model-based* indices of control adjustment costs).

#### Model-Agnostic Evidence of Control Adjustment Costs

As shown in Figure 2A and Table S1, participants were slower (*b*=128.25; 95% CrI [85.36, 172.52]; *p_b_*_<0_=0.00) and more accurate^1^ (*b*=1.67; 95% CrI [1.12, 2.22]; *p_b_*_<0_=0.00) when their performance goal was to maximize accuracy compared to when it was to maximize speed. Crucially, we found evidence for control adjustment costs when comparing fixed and varying blocks. The difference between speed and accuracy conditions was larger in fixed compared to varying blocks for both reaction times (interaction effect: *b*=28.09; 95% CrI [3.54, 51.41]; *p_b_*_<0_=0.01) and accuracy (interaction effect: *b*=0.44; 95% CrI [0.12, 0.76]; *p_b_*_<0_=0.00). These results indicate that the performance in the speed condition was closer to the performance in the accuracy condition when participants had to switch between these strategies within one block relative to when they had only one performance goal per block. We confirmed (Table S2) that these findings are not driven by changes in vigor (non-decision time) nor the reduction of thresholds over time (collapsing bound).

We next analyzed behavior in the varying block, focusing our analysis on whether the performance goal on the previous interval was the same as on the current one (repeat), or different than the current goal (switch). This analysis (Figure 2C) revealed that the difference between accuracy and speed performance goals (reaction times: *b*=113.19; 95% CrI [71.51, 156.99]; *p_b_*_<0_=0.01; accuracy: *b*=1.45; 95% CrI [0.87, 2.02]; *p_b_*_<0_=0.01), was reduced when the current interval included a goal switch compared to a goal repeat (interaction effects; reaction times: *b*=10.97; 95% CrI [-1.65, 23.86]; *p_b_*_<0_=0.04; accuracy: *b*=0.23; 95% CrI [0.01, 0.46]; *p_b_*_<0_=0.02).

#### Model-based Evidence of Control Adjustment Costs

Our model predicts that, due to control adjustment costs, drift rates and thresholds for the two performance goals will be closer together in blocks in which the goals vary compared to blocks in which they are fixed (Figure 2B-left). To test this prediction, we fitted a drift diffusion model to behavioral data in order to jointly model both the reaction times and accuracies, and to directly probe the prediction from our model. This analysis revealed a substantial difference in both drift rates and thresholds between the speed and accuracy performance goals. Participants had higher drift rates (*b*=0.54; 95% CrI [0.31, 0.77]; *p_b_*_<0_=0.00) and thresholds (*b*=0.42; 95% CrI [0.27, 0.58]; *p_b_*_<0_=0.00) when they were instructed to focus on accuracy, relative to when they were instructed to focus on speed (Figure 2B-right and Table S1).

As predicted by our model (Figure 2B-left), the distance between the speed and accuracy conditions was reduced in varying compared to fixed blocks. This was true in thresholds (interaction effect: *b*=0.08; 95% CrI [0.03, 0.13]; *p_b_*_<0_=0.00; Figure 2B-right), and held in the same direction for drift rates, but not reliably so (interaction effect: *b*=0.05; 95% CrI [-0.11, 0.21]; *p_b_*_<0_=0.28). To obtain a single metric of control adjustment costs, we computed Euclidian distances between the speed and accuracy conditions within the two-dimensional space formed by the values of drift rate and threshold (see Methods for details). This analysis showed that the distance between the two conditions was reduced in varying compared to fixed blocks (*b*=0.11; 95% CrI [-0.01, 0.24]; *p_b_*_<0_=0.04; Figure 6), indicating the existence of control adjustment costs.

The second prediction from our model is that the distance between drift rates and thresholds among the speed and accuracy conditions should be reduced on switch compared to repeat intervals (Figure 2D-left and Table S3). To test this prediction, we focused our analysis on the varying blocks and compared the difference between accuracy and speed intervals in cases which required a switch to the cases which did not require switches (i.e., repeats of the performance goal). This analysis revealed that the effect of performance goal on thresholds was larger in repeat relative to switch intervals (interaction effect: *b*=0.04; 95% CrI [0.00, 0.07]; *p_b_*_<0_=0.02). Similar to the block-wise analysis, this difference was not found for the drift rates (interaction effect: *b*=0.00; 95% CrI [-0.15, 0.15]; *p_b_*_<0_=0.49). These findings closely follow the block-wise analyses, suggesting that the differences in control intensities between fixed and varying blocks arise due to the control adjustment costs stemming from the need to frequently adjust control signals in response to changing performance goals.

For completeness, we tested the role of Stroop conflict between targets (ink color) and distractors (color word). We observed the typical congruency effect, participants were slower to respond (*b*=39.18; 95% CrI [28.82, 49.61]; *p_b_*_<0_=0.00) and less accurate (*b*=-0.25; 95% CrI [-0.36, -0.14]; *p_b_*_>0_=0.00) on incongruent compared to congruent trials. Furthermore, we found that control adjustment costs (larger difference between speed and accuracy conditions in fixed compared to varying blocks) were present in both congruent and incongruent trials (Table S4; Figure S1). Like in our previous work (Leng et al., 2021), congruency effect was captured within the Drift Diffusion model through its influence on the starting point bias and drift rate parameters. Compared to congruent trials, incongruent trials demonstrated a bias toward the incorrect response boundary (*b*=-0.02; 95% CrI [-0.03, -0.01]; *p_b_*_>0_=0.00) and a lower drift rate (*b*=-0.34; 95% CrI [-0.41, -0.28]; *p_b_*_>0_=0.00). These effects of congruency on reaction times, accuracy, and Drift Diffusion model parameters were observed across all 4 studies (Supplementary materials).

#### Discussion

The results of Study 1 confirmed that we can induce reliable differences in behavioral performance and in estimates of underlying control signal intensities (drift rates and thresholds). This allowed us to test for the existence of control adjustment costs predicted by our model. Comparing environments which demand frequent changes in goals states to the ones which don’t, we found evidence for control adjustment costs in both behavioral performance and control intensity. Confirming the first prediction of our model, we found that control adjustment costs arise due to the demand to switch between different performance goals. Next, we tested the prediction of our model that control adjustment costs are governed by the distance between control states, with longer distances producing larger control adjustment costs.

### The Costs of Control Adjustment Scale with the Distance Between Control States (Study 2)

Our model assumes that control adjustment costs depend on the distance between the current state of the control system and the desired state induced by the new performance goal. The model assumes that control intensities are gradually adjusted toward the desired state. A key prediction that follows is that the distance between the current and desired state determines control adjustment costs. We should therefore observe greater adjustment costs (as indexed by the distance between target control levels and implemented control levels) the further the current goal state is from the previous one. Study 2 tested this prediction by having participants (N = 38) perform the Stroop task under three performance goals: respond as quickly as possible (speed goal), as accurately as possible (accuracy goal), or both as fast and accurately as possible (speed/accuracy goal). Participants performed the task in blocks in which the goal was always the same (fixed blocks), or in blocks in which the three goals randomly changed (varying blocks). This paradigm allowed us to investigate whether control adjustment costs depend on the distances between different target goal states.

#### Model Predictions

Simulations of this task were generated in the same way as in Study 1. We first simulated fixed blocks in which the agent performed under only one of the three performance goals (speed, accuracy, and speed/accuracy). We then compared performance in fixed blocks to simulated performance in varying blocks in which the agent switched between the three performance goals. The model predicted (Figure 3B-left) larger control adjustment costs for the conditions that were further in control state space (speed vs. accuracy), relative to the conditions which were closer (accuracy vs. speed/accuracy).

**Figure 3.**
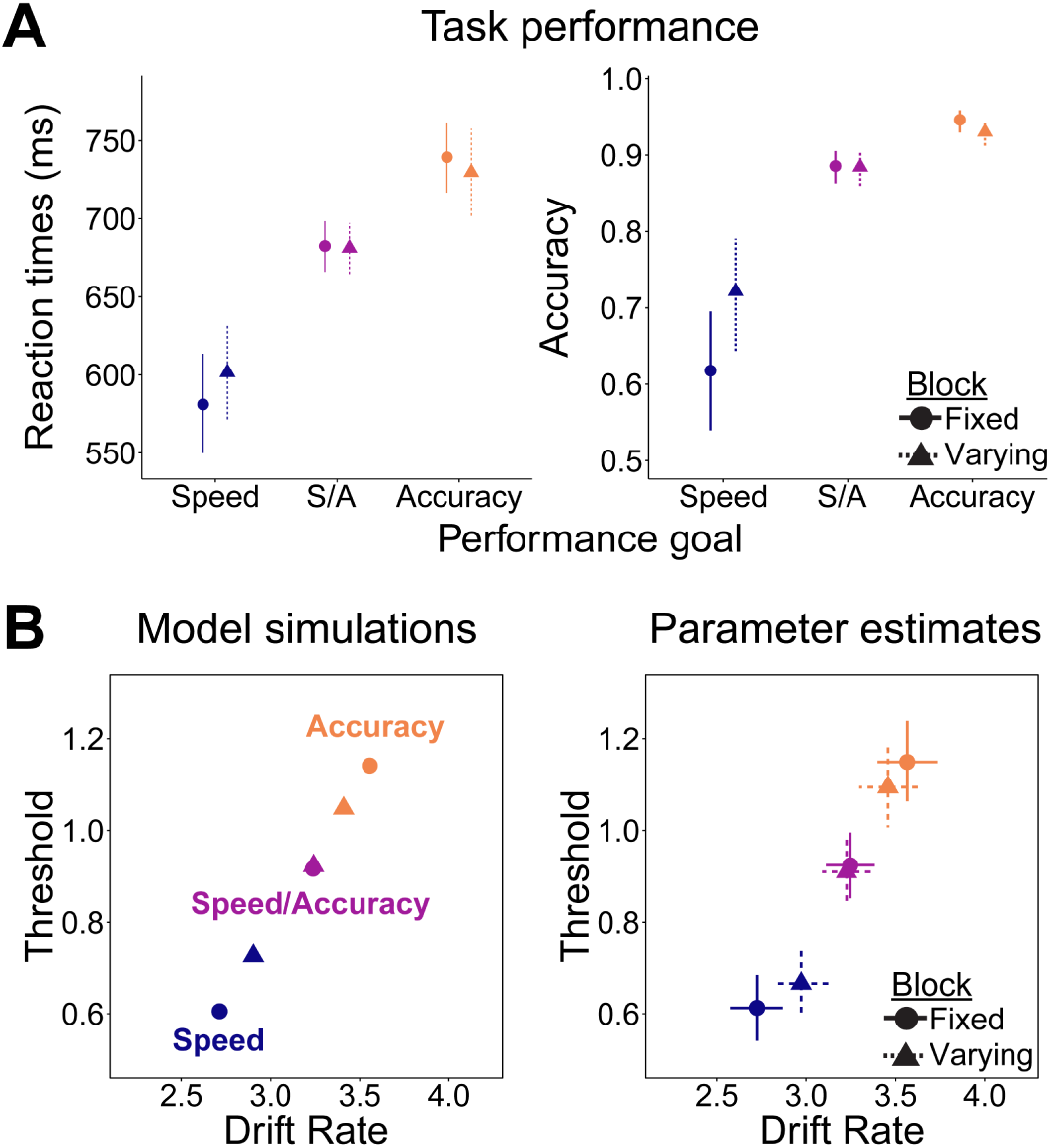
Magnitude of control adjustments impacts behavioral performance and control intensity. **A.** Reaction times and accuracies for the three performance goals in fixed and varying blocks. Dots represent regression estimates and error bars 95% credible intervals. **B.** Model simulations (left) and drift diffusion parameter estimates (right) for the three performance goals. Error bars represent 95% credible intervals.

#### Model-Agnostic Evidence of Control Adjustment Costs

Replicating the first experiment, we found that participants were slower (*b*=143.32; 95% CrI [95.18, 190.48]; *p_b_*_<0_=0.00) and more accurate (*b*=2.01; 95% CrI [1.50, 2.52]; *p_b_*_<0_=0.00) when they performed under the accuracy compared to the speed goal (Figure 3A and Table S5). In the speed/accuracy condition, participants’ performance was in between that of the other two performance goals. Relative to the accuracy condition, participants were faster (*b*=-52.74; 95% CrI [-79.68, -26.12]; *p_b_*_>0_=0.00), but less accurate (*b*=-0.69; 95% CrI [-79.10, -25.10]; *p_b_*_>0_=0.00). Thus, these three performance goals induced distinct behavioral performance profiles.

Replicating Study 1, we found evidence of control adjustment cost when comparing speed and accuracy conditions in varying and fixed blocks. The costs were present in both reaction times (interaction effect: *b*=30.30; 95% CrI [-3.83, 64.80]; *p_b_*_<0_=0.04) and accuracy (interaction effect: *b*=0.75; 95% CrI [0.31, 1.20]; *p_b_*_<0_=0.00). Critically, this control adjustment costs (i.e., performance difference between performance goals across fixed and varying blocks) was not found when comparing other performance goals which were closer together, thus requiring less adjustment. We found no evidence of adjustment costs between accuracy and speed/accuracy conditions (interaction effects; reaction times: *b*=-8.43; 95% CrI [-35.40, 17.49]; *p_b_*_>0_=0.26; accuracy: *b*=-0.27; 95% CrI [-0.64, 0.10]; *p_b_*_>0_=0.08), nor between speed and speed/accuracy (interaction effects; reaction times: *b*=-26.65; 95% CrI [-61.21, 8.64]; *p_b_*_>0_=0.10; accuracy: *b*=-0.07; 95% CrI [-0.39, 0.25]; *p_b_*_>0_=0.36).

#### Model-based Evidence of Control Adjustment Costs

During fixed blocks, participants had the highest drift rates and thresholds in the accuracy condition, and lowest in the speed condition (Figure 3B-right). The speed/accuracy condition was in between the other two. We found significant differences between both accuracy and speed/accuracy conditions (drift rates: *b*=0.27; 95% CrI [0.09, 0.46]; *p_b_*_<0_=0.01; thresholds: *b*=0.20; 95% CrI [0.10, 0.31]; *p_b_*_<0_=0.00) and between speed/accuracy and speed conditions (drift rates: *b*=0.39; 95% CrI [0.27, 0.51]; *p_b_*_<0_=0.00; thresholds: *b*=0.28; 95% CrI [0.20, 0.35]; *p_b_*_<0_=0.00). The distance between the speed and the speed/accuracy condition was larger than the distance between the accuracy and the speed/accuracy condition, but this difference was not robust (drift rates: *b*=0.21; 95% CrI [-0.10, 0.51]; *p_b_*_<0_=0.13; thresholds: *b*=0.08; 95% CrI [-0.07, 0.24]; *p_b_*_<0_=0.18). Thus, the three performance goals successfully produced different control states. This allowed us to test the prediction from our model that conditions farther away in control space (speed vs. accuracy) should result in higher control adjustment costs than conditions closer together (speed/accuracy vs. accuracy and speed/accuracy vs. speed).

Replicating Study 1, we found evidence of control adjustment costs when comparing the differences between the accuracy and speed goals in fixed and varying blocks (interaction effects; drift rates: *b*=0.36; 95% CrI [0.16, 0.55]; *p_b_*_<0_=0.00; thresholds: *b*=0.11; 95% CrI [0.04, 0.18]; *p_b_*_<0_=0.00; Figure 3B-right and Table S5). Of note, in this experiment we found adjustment costs in both threshold and in drift rate (which was not significant in Study 1). Critically, we confirmed the second prediction of our model, as we found no evidence of control adjustment costs between two closest conditions – accuracy and speed/accuracy – in neither drift rates (interaction effect; *b*=0.09; 95% CrI [-0.09, 0.27]; *p_b_*_<0_=0.21), nor thresholds (interaction effect; *b*=0.04; 95% CrI [-0.03, 0.11]; *p_b_*_<0_=0.17). However, we did find significant adjustment costs, though smaller in magnitude, when comparing the speed/accuracy to the speed condition (interaction effects; drift rates: *b*=0.27; 95% CrI [0.03, 0.51]; *p_b_*_<0_=0.03; thresholds: *b*=0.07; 95% CrI [0.02, 0.12]; *p_b_*_<0_=0.01).

These findings were confirmed when applying our Euclidean distance-based estimate of adjustment costs (i.e., the distance between control configurations in fixed versus varying block). These cost estimates were greater when goal states were further apart (accuracy vs. speed: *b*=0.24; 95% CrI [0.11, 0.38]; *p_b_*_<0_=0.00, speed/accuracy vs. speed: *b*=0.17; 95% CrI [0.03, 0.31]; *p_b_*_<0_=0.00) but not when goal states were closer together (accuracy vs. speed/accuracy: *b*=0.07; 95% CrI [-0.05, 0.20]; *p_b_*_<0_=0.14) (Figure 6). Thus, we found that distance in control space determines control adjustment costs, with increased costs in situations that demand larger adjustments in control.

#### Discussion

Results of Study 2 confirmed our model’s prediction that control adjustment costs depend on the distance between the current and the desired goal state. By investigating behavioral performance and control signal intensities across three performance goals we showed that the costs increase as a function of the distance between the goal-optimal points in the control state space. We next tested the prediction that control states are gradually adjusted over time. We thus tested the impact of time allowed to change control signals on control adjustment costs.

### The Costs of Control Adjustments Depend on the Time Allowed for Adjustments (Study 3)

Our model assumes that control signals are adjusted incrementally from the current to the desired control state. To investigate the role of time in control adjustments, in this study we had participants (N=50) perform the same task as in Study 1 (speed vs. accuracy goals in fixed and varying blocks), but we varied the amount of time they had to adjust control signals. In half of the blocks participants had a shorter amount of time to adjust control (250ms stimulus-onset asynchrony (SOA) between the goal cue and task onset), while in the other half of blocks they had more time (1000ms SOA) to adjust control. In addition, in this experiment we made goal switches fully predictable (switch every interval) to ensure that control adjustment costs are not dependent on whether on uncertainty about the current or the next goal.

#### Model Predictions

Simulations of this study were the same as in Study 1, but we now also simulated blocks with shorter or longer SOA by having the model take either 250 or 1000 steps before implementing the reached control state. The model predicts that shorter adjustment windows will result in increased control adjustment costs because there is less time to move toward the desired control state before task onset (Figure 4B-left).

**Figure 4.**
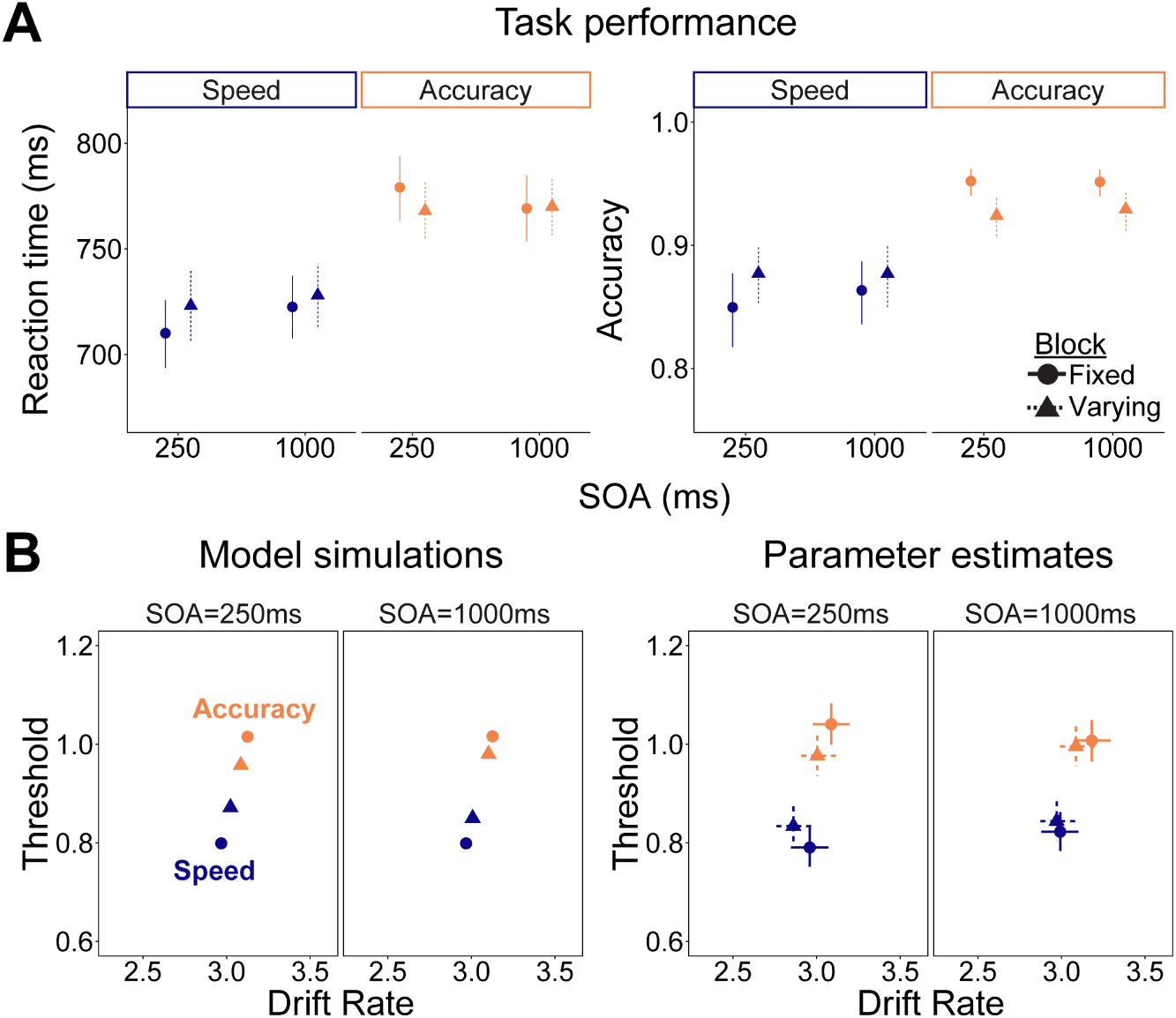
Control adjustment costs are increased when there is less time to change control signals. **A.** Reaction times and accuracies for the speed and accuracy performance goals when the time to adjust control (SOA between goal cue and task onset) is 250 vs. 1000ms. Dots represent regression estimates and error bars 95% credible intervals. **B.** Model simulations (left) and drift diffusion parameter estimates (right) speed and accuracy performance goals when SOA is 250 vs. 1000ms. Error bars represent 95% credible intervals.

#### Model-Agnostic Evidence of Control Adjustment Costs

Replicating our previous studies, we found that participants were slower (*b*=50.66; 95% CrI [29.08, 71.58]; *p_b_*_<0_=0.00) and more accurate (*b*=0.88; 95% CrI [0.57, 1.20]; *p_b_*_<0_=0.00) when instructed to focus on accuracy relative to speed (Figure 4A and Table S6). We again found evidence for control adjustment costs when comparing performance in varying and fixed blocks (interaction effects; reaction times: *b*=14.37; 95% CrI [-1.68, 29.37]; *p_b_*_<0_=0.04; accuracy: *b*=0.62; 95% CrI [0.36, 0.87]; *p_b_*_<0_=0.00). We found suggestive evidence indicating that control adjustment costs were larger in blocks in which people had shorter time to adjust (SOA =250ms) compared to when they had more time (SOA=1000ms) in both reaction times (interaction effect; *b*=-19.55; 95% CrI [-51.10, 11.65]; *p_b_*_>0_=0.10) and accuracy (interaction effect; *b*=-0.20; 95% CrI [-0.61, 0.19]; *p_b_*_>0_=0.16). While these interactions were not individually significant, both RT and accuracy effects were directionally consistent with the expected direction of change, suggesting that they may jointly reflect a reliable change in the relevant model-based indices of control costs.

#### Model-based Evidence of Control Adjustment Costs

Replicating our previous results, we found (Figure 4B-right and Table S6) that drift rates (*b*=0.14; 95% CrI [0.06, 0.23]; *p_b_*_<0_=0.00) and thresholds (b=0.18; 95% CrI [0.12, 0.24]; *p_b_*_<0_=0.00) were higher in the accuracy relative to speed condition. We also found evidence for control adjustment costs in threshold (interaction effect; *b*=0.07; 95% CrI [0.03, 0.11]; *p_b_*_<0_=0.00). Critically, control adjustment costs were reduced when participants had more time to adjust control (interaction effect; *b*=0.07; 95% CrI [0.00, 0.15]; *p_b_*_>0_=0.03). For drift rates, we found no evidence of control adjustment cost, nor a change in the cost due to the amount of time to adjust control (*p*s>0.3). This finding was recapitulated in our Euclidean metric of control adjustment costs, measuring change in distances between goal-specific control configurations between fixed and varying blocks. Using this metric, we found a significant cost (change in distance) when participants had 250ms to adjust control (*b*=0.24; 95% CrI [0.06, 0.41]; *p_b_*_<0_=0.00), but the cost was smaller and not reliably above zero when they had 1000ms to adjust control (*b*=0.11; 95% CrI [-0.07, 0.30]; *p_b_*_<0_=0.11). While this change in control cost between the conditions with more or less time to adjust was not significant (*b*=0.13; 95% CrI [-0.14, 0.37]; *p_b_*_<0_=0.11), it was in the direction predicted by our model. These results confirm our model’s prediction that suboptimal adjustments can arise from insufficient time to transition between current and target levels of control (Figure 4B-left).

#### Discussion

In this study we demonstrated that control adjustment costs depend on the time allowed for the adjustment. Since our model assumes that control states are gradually adjusted from the current toward the desired state, the model predicts decreases in adjustment costs as the time allowed for the adjustment increases. Our findings confirm this prediction, showing that control adjustment costs were higher when participants had less time to adjust control. Next, we tested the prediction that control adjustment costs depend on the frequency of goal switches, with more frequent switches producing larger adjustment costs.

### Control Adjustment Costs Depend on Goal Switch Frequency (Study 4)

Our model predicts that the number of goal state switches determines the control adjustment costs. In this experiment, we directly tested this prediction by comparing fixed blocks (no goal switches) to varying blocks (randomly in Studies 1 and 2 and every interval in Study 3). Further, we explicitly instructed participants about the goal switch frequency prior to the start of each block (cf. Liu & Yeung, 2020). Thus, we could investigate the effect of goal switch frequency on control adjustment costs. Participants (N=55) performed the same Stroop task under speed and accuracy goals. Again, we had fixed blocks in which the goal never changed. In three other block types, goals predictably switched with low (every 8 intervals), medium (every 4 intervals), or high frequency (every 2 intervals).

#### Model Predictions

To generate model predictions for this study, we simulated the conditions in which the speed and accuracy performance goals never change (fixed blocks), as well as the blocks in which the goals change with low, medium, or high frequency (Figure 5B). The model predicts that control adjustment costs increase with increasing frequency of goal switches. Following a goal change, the simulated agent will slowly adjust their control state (e.g., several trials). When switch frequency is low, the agent will reach their target control state and then perform a few more trials in that state. However, when switch frequency is high, the agent will barely reach the state before having to adjust it again due to a new goal change.

**Figure 5.**
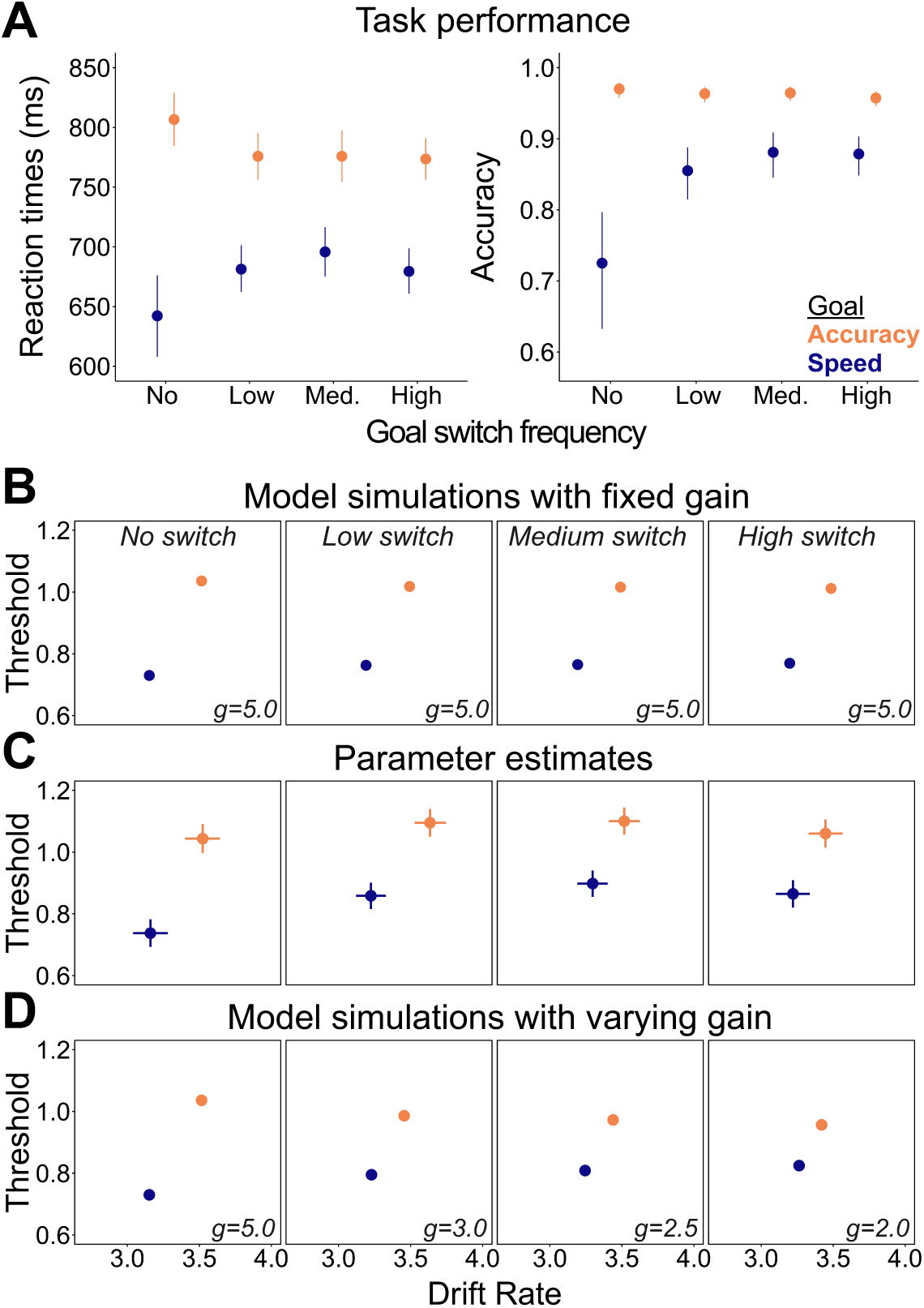
Control adjustment costs increase with the number of performance goal switches. **A.** Reaction times and accuracies across blocks in which performance goals are fixed (no switches), or they change every 8 (low), 4 (medium), or 2 (high) intervals. Dots represent regression estimates and error bars 95% credible intervals. **B.** Model simulations across four levels of goal switch frequency. This model predicts increasing control adjustment costs with the increase of the goal switch frequency. **C.** Drift diffusion parameter estimates. Error bars represent 95% credible intervals. **D.** Simulations from the model which assumes that participants proactively adjust gain based on the instructions informing them about the goal switch frequency within a block. This model captures the magnitude of the empirical effect better than the model with fixed gain.

**Figure 6.**
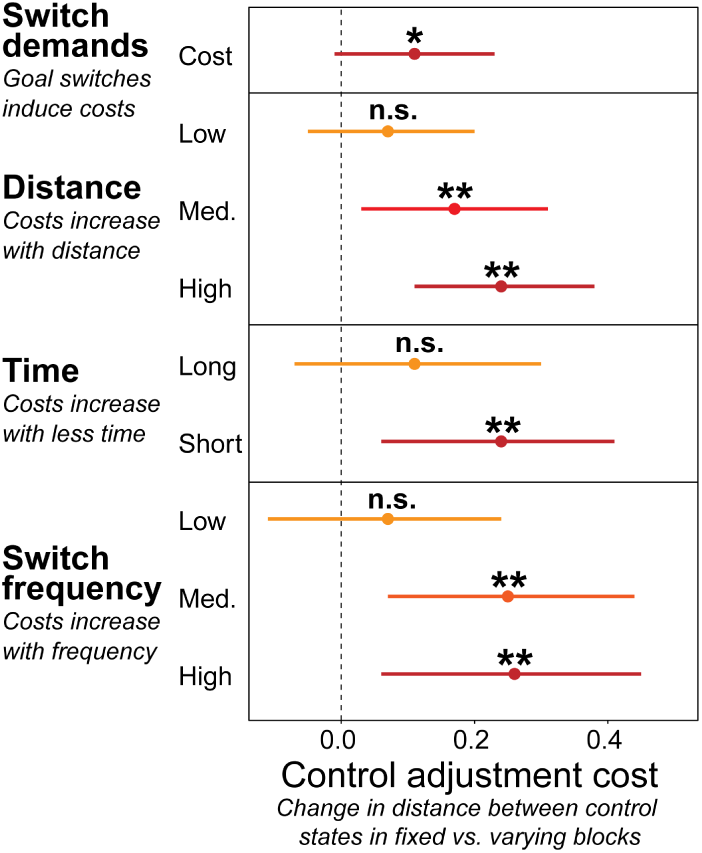
Summary of model-based control adjustment costs and their causes across four studies. We calculated Euclidian distances between control states (drift rates and thresholds) corresponding to the speed and accuracy goals. Control adjustment costs are indexed by the reduction in this distance in blocks that require frequent control adjustments (varying blocks) relative to blocks with no adjustments (fixed blocks). Points represent means of the posterior distributions obtained from Drift Diffusion Model estimates, and error bars are 95% credible intervals. Study 1 shows a significant adjustment cost on threshold. Study 2 shows the smallest cost between states with low distance (Accuracy vs. Speed/Accuracy goal), larger cost with medium distance (Speed/Accuracy vs. Speed), and largest cost with high distance (Accuracy vs. Speed goal). Study 3 shows a larger cost in threshold when people have less time to adjust control states. Study 4 shows that relative to the fixed block, the cost is smallest when people switch goals every 8 intervals, larger when they switch every 4 intervals, and largest when they switch every 2 intervals.

#### Model-Agnostic Evidence of Control Adjustment Costs

Participants were once again slower (*b*=54.14; 95% CrI [37.96, 70.30]; *p_b_*_<0_=0.00) and more accurate (*b*=0.80; 95% CrI [0.58, 1.04]; *p_b_*_<0_=0.00) when instructed to focus on accuracy compared to speed (Figure 5A). Critically, control adjustment costs (goal type vs. block interaction) increased with the increasing frequency of goal switches (Figure 5A and Table S7). This was true for both the reaction times (goal vs. block vs. switch frequency interaction effect; *b*=25.20; 95% CrI [8.50, 42.58]; *p_b_*_<0_=0.00) and accuracy (goal vs. block vs. switch frequency interaction effect; *b*=0.48; 95% CrI [0.25, 0.74]; *p_b_*_<0_=0.00). This result suggested that control adjustment costs depend on the number of times participants had to adjust control intensity.

#### Model-based Evidence of Control Adjustment Costs

Replicating our previous studies, we found higher drift rates (*b*=0.30; 95% CrI [0.17, 0.44]; *p_b<0_*=0.00) and thresholds (*b*=0.23; 95% CrI [0.17, 0.29]; *p_b<0_*=0.00) in accuracy compared to speed intervals. Importantly, as predicted, control adjustment costs increased with more goals switches in a block (Figure 5C and Table S7), for both drift rates and thresholds (goal vs. block vs. switch frequency interaction effect; drift rates: *b*=0.20; 95% CrI [0.05, 0.43]; *p_b>0_*=0.06; thresholds: *b*=0.12; 95% CrI [0.04, 0.21]; *p_b<0_*=0.00). As in our earlier studies, these findings were recapitulated in our Euclidean metric, which identified significant control adjustment costs when goals changed with high frequency (*b*=0.26; 95% CrI [0.06, 0.45]; *p_b_*_<0_=0.01), or medium frequency (*b*=0.25; 95% CrI [0.07, 0.44]; *p_b_*_<0_=0.00), but not when they changed with low frequency (*b*=0.07; 95% CrI [-0.11, 0.24]; *p_b_*_<0_=0.23), relative to fixed blocks. Thus, control adjustment costs varied with the expected frequency of goal switches, confirming the predictions generated from the control dynamics model (Figure 5B).

#### A Role for Dynamic Gain Adjustments

Our model predicts that control states will be pressed closer together (i.e., increased adjustment costs) as goal switch frequency increases (Figure 5B). While this prediction is confirmed by our data, the magnitude of the observed effect is substantially larger than predicted (Figure 5C). Importantly, unlike in the other studies reported above, participants in this study were provided with advance information about goal switch frequency in the upcoming block. Thus, we ran an additional set of simulations to test whether proactive adjustments in cognitive flexibility could explain the mismatch between the predicted and the observed magnitude of control adjustment costs across levels of goal switch frequency.

Previous work on task-switching has shown that frequent switches demand higher flexibility, which can be captured in models with changing how sensitive the model is to the input about the current task (Ueltzhöffer et al., 2015; Musslick et al., 2018; Musslick et al., 2019). Lowering the sensitivity to task inputs allows for more flexible control configurations, and thus, is optimal in environments in which task goals frequently switch. However, such flexible control configurations come at a cost: they degrade the performance of each individual task (Musslick & Bizyaeva, 2024; Musslick & Cohen, 2021). To implement this in our model we adjusted the gain parameter (*g* in Equation 1) of the sigmoid activation function across the 4 block types (see Methods for details). Thus, g can be thought of as a meta-parameter, that may be set by the participant depending on their expectation about the volatility of the environment. Following previous work (Musslick, Bizyaeva, et al., 2019; Musslick et al., 2018), we assumed that participants could proactively adjust gain. We parametrically decreased gain across blocks with increasing number of goal switches. This pressed the control configurations closer together, effectively reducing the distance that participants had to travel between control states. The model that included such a proactive manipulation of flexibility captured the observed data better than the model with fixed gain (Figure 5D).

#### Discussion

In this study we confirmed that control adjustment costs depend on the frequency with which participants have to adjust their control states. We show that the costs parametrically increase with the number of goal switches within a block. This finding validates the prediction that arises from the control dynamics model we developed, further validating the proposal that control adjustment costs arise due to the time needed to adjust control states when performance goals change. Furthermore, we provide initial evidence showing that people can proactively adjust how flexible they are based on expectations about goal switch demands. The empirically observed data are better captured by a model that assumes that people increase flexibility when entering blocks with frequent goal changes. This flexibility adjustment effectively presses their control states closer together, allowing for more rapid switches, but at the cost of being further away from the ideal target control state.

### Alternative Models

We have proposed a model in which control states are gradually adjusted from their current state to the target state imposed by a new performance goal. However, other models could make similar predictions. Here we lay out those predictions, and then directly compare them to our model.

#### Control States are Instantaneously Adjusted

Recent computational models have modeled task switching as a gradual activation of a new task set and a gradual deactivation of the previous task set (e.g., Ueltzhöffer et al., 2015; Musslick et al., 2018; Steyvers et al., 2019). Following this work, we have proposed a model in which control states are gradually adjusted following a goal change, even in absence of a task set change. An alternative to this is that cognitive flexibility is limited only by the costs of switching between task sets, but that there are no additional costs to adjusting control states that govern how quickly or accurately people perform a task. Such a model would postulate that people immediately “jump” between control states once their performance goal changes. This model would predict no change in distance between control states when comparing fixed to varying blocks. To directly compare this model to the model we proposed, we fit two Drift Diffusion Models to the data in Study 1. Both models had all the predictors previously outlined, but one model included control adjustment costs (i.e., the interaction between block type and goal type), while the other model didn’t (i.e., only the main effects of block type and goal type). This analysis revealed that the model including the costs (DIC = -1918) outperforms the model without the costs (DIC = -1745).

#### Adjusting Control States Incurs a Fixed Cost

While the previous model comparison demonstrates that a model without control adjustment costs doesn’t fit the data well, this comparison does not reveal anything about the cause of these costs. The model we have developed proposes that adjustment costs arise due to gradual adjustments of control states. Critically, the model states that control dynamics are governed by the distance between the current and the new target control state. However, a plausible alternative would be that there is a fixed control adjustment cost. Such a model predicts the existence of a cost (i.e., interaction between block type and goal type), but does not predict that the magnitude of the cost is modulated by the time allowed to adjust control. Data collected in Study 3 allow us to directly compare our model to the model with a fixed control adjustment cost. Model comparison showed that the model which includes the modulation of costs by time outperforms (DIC = -1918) the model that includes the cost, but only the main effect of time (DIC = -1745).

## General Discussion

One of the hallmarks of human behavior is the ability to flexibly adjust to the demands of the current environment. Frequent changes in our goals, as well as in our environments, demand adjustments in our behavior. These adjustments are supported by changing cognitive control states, but such changes incur costs. Previous work has focused on the costs that arise from the need to adjust discrete control states determining which stimuli are processed and which actions are taken (Allport et al., 1994; Rogers & Monsell, 1995; Monsell, 2003; Kiesel et al., 2010). Here we extend the understanding of the costs of switching between goals by focusing on adjustment in control signals that determine levels of attention and response caution. We show that adjustments in cognitive control levels produce robust costs in environments that demand frequent goal switches. We expose these control adjustment costs in a novel behavioral paradigm and test predictions of a computational model showing that the costs arise due to the dynamics of changes in cognitive control states that support goal-directed behavior.

Here we focus on adjustments of continuous control states defined by the amount of attention and caution required by the current goal. Such control states broaden the notion of a task-set, which commonly specifies a control state with respect to which stimuli are attended and which responses are taken (Allport et al., 1994; Rogers & Monsell, 1995). Mounting empirical evidence shows that people adjust their performance within a task based on performance goals (Forstmann et al., 2008; Ratcliff & Rouder, 1998) and expected incentives (Manohar et al., 2015; Parro et al., 2018; Frömer et al., 2021). The Drift Diffusion model (Bogacz et al., 2006; Ratcliff & McKoon, 2008; Ritz et al., 2022) is commonly used to measure changes in response speed and accuracy as a function of instructions or incentives (Forstmann et al., 2008; Leng et al., 2021; Grahek et al., 2023). Here we leverage this model to conceptualize control states along a two-dimensional space, defined by levels of attention and response caution demanded by the current goal (e.g., try to perform a task quickly or accurately). This allowed us to study continuous adjustments of cognitive control states, defined by *how efficiently* people processed stimuli (drift rate parameter, a putative analog of top-down attention) and *how cautious* they are when they respond (threshold parameter). We showed that costs arise from the requirement to transition between the target control state for one goal (e.g., low attention and low caution) and the target control state for a different goal (e.g., high attention and high caution) (Study 1). We show that such costs arise in particular from a combination of the distance one needs to traverse between control states (Study 2) and the time allowed for that transition process (Study 3), with greater costs (smaller adjustments) emerging when target control states are far apart (e.g., Speed goal → Accuracy goal) and/or transition time is short (e.g., 250ms), relative to when those control states are closer (e.g., Speed → Speed & Accuracy) and the transition time is longer (e.g., 1000ms) and subject to inertia. We show that these costs also increase as the goal switches become more frequent, and that people can proactively use the information about goals switch frequency to adjust how flexible they are within an environment (Study 4). This work offers a mechanistic account of the causes of costs associated with adjusting cognitive control states.

Computational models of cognitive control posit that people adapt their allocation of control to a task (including levels of attention and caution) based on the expected benefits and costs of doing so. Across different models, optimal control states are found by maximizing the expected utility of control signals (Bogacz et al., 2006; Simen et al., 2006; Shenhav et al., 2013; Manohar et al., 2015; Verguts et al., 2015; Alexander et al., 2018). In support of these models, empirical work shows that people adjust control according to their current task goal (e.g., focus on speed vs. accuracy; Forstmann et al., 2008; Ratcliff & Rouder, 1998), expected rewards (Padmala & Pessoa, 2011; Krebs et al., 2012) and penalties (Leng et al., 2021), and based on the expected costs of control (Kool et al., 2010; Westbrook et al., 2013, 2020). Thus, these models assume that levels of control depend on the incentives in one’s environment, and that these will be adjusted as incentives change (either extrinsic incentives or goals). While some of the computational models incorporate a cost for adjusting control from one setting to another (Musslick et al., 2015; Lieder et al., 2018; Grahek et al., 2020), such costs were until now only stipulated and not empirically demonstrated. Our work provides empirical support for these models, but also goes further by specifying how these costs emerge.

Investing higher levels of control (e.g., allocating more top-down attention to task-relevant stimuli) and switching between goals both incur costs. While the causes of both types of costs remain deeply mysterious (Shenhav et al., 2017), an intriguing possibility is that these two costs are linked, with the latter in part explaining the former. Specifically, drawing on past research on the stability-flexibility dilemma – the observed trade-off between being able to focus cognitive resources on the task at hand versus rapidly re-adjust information processing (Goschke, 2013) – Musslick & Cohen (2021) propose that the control system may have a bias against maximizing its allocation of control resource to a given task because this would make it more difficult to rapidly switch to a new task. Their account predicts that the allocation of control towards a given task depends in part on the expectation about the frequency of task switches in the current environment (Musslick, Bizyaeva, et al., 2019). Our theoretical and empirical findings give substance to this account. First, we propose that control adjustment costs arise from the time it takes to adjust control states from their current to the reward-optimal states. Second, we show that our model can be cast as a neural network, and that the balance between stability and flexibility can be set by adjusting the gain of the activation function (cf., Ueltzhöffer et al., 2015). We show that changes in the gain of the activation function that correspond to the expected frequency of control state switches are able to capture our empirical data well (Study 4). Finally, and of particular note, our model is able to capture these control adjustment costs without explicitly incorporating them into the evaluation that determines control allocation (an assumption that has been incorporated into some previous models in order to account for switch costs; Musslick et al., 2015; Lieder et al., 2018; Grahek et al., 2020). Rather, we propose that control adjustment costs arise from the time it takes to adjust control states, and the distance between the current and the desired control state. In this sense, control adjustment costs arise due to the control dynamics inherent in the configuration of the cognitive/neural system, and thus do not require further assumptions about the root causes of the cost. This proposal aligns well with the recent focus on the dynamics of control (Kikumoto & Mayr, 2020; Steyvers et al., 2019; Holroyd, 2023), as well as with proposals that cognitive effort scales with how hard it is to reach the desired control states from the current states (Kim et al., 2018; Weninger et al., 2022).

The current study prompts a critical inquiry: do the costs that we report here derive from the same mechanisms postulated for inter-task switches? Similar to commonly observed switch costs (Kiesel et al., 2010), we find performance differences between trials involving goal switches and goal repetitions (i.e., local costs), and blocks with and without goal switches (i.e., mixing costs) within the same task. However, unlike the classical switch costs, our results show that performance is not simply degraded on switch trials and varying blocks, but is rather pulled toward the previously implemented control state due to inertia in control adjustments. Nevertheless, the core mechanisms causing both control adjustment costs and task switch costs could be similar. Some researchers advocate that task switch costs emanate from an active task-set reconfiguration process, underpinned by a control mechanism (Rogers & Monsell, 1995; Meiran, 1996; Mayr & Kliegl, 2000). In contrast, others propose these costs stem from passive processes such as proactive interference — also termed “task-set inertia” (Allport et al., 1994), inhibition of the last performed task-set (Mayr & Keele, 2000; Altmann, 2007), repetition priming of the task cue (Logan & Bundesen, 2003), or repetition priming of stimulus attributes (Wylie & Allport, 2000; Waszak et al., 2004). Our findings align with the passive process theory, ascribing within-task switch costs to the inertia from reconfiguring control states. In this sense, they provide further evidence for Allport’s task-set inertia hypothesis, extend it to the domain of continuous control states, and in doing so provide a way to directly measure inertia. Nevertheless, these results introduce further queries, notably whether within-task control reconfiguration costs can be tied to other passive mechanisms like repetition priming, or perhaps to active control state reconfiguration processes. A promising avenue for tackling these questions would be to investigate whether control dynamics, and the ensuing adjustment costs, can be proactively modulated by incentivizing fast control adjustments. Furthermore, potential differences in dynamics either within (e.g., time on task) or between individuals (e.g., aging) could elucidate the extent to which control dynamics can be modulated.

Our ability to flexibly adjust control processes lies at the core of goal-directed behavior. Given the importance of such behavior, the question of why there are costs to adjusting cognitive control state which support it, remains a puzzle within cognitive science (Musslick & Masís, 2023). By focusing on adjustments of continuous control states, rather than changes in discrete task-sets, we show how costs of adjusting control can arise from the dynamics of cognitive control. Many real-world tasks include such adjustments in attention and caution, but this aspect of control adjustments is often overlooked. Thus, studying control adjustment costs opens an avenue for a more precise understanding of the mechanisms which lead to behavioral inflexibility, which is critical for understanding both psychopathology (Kashdan & Rottenberg, 2010) and aging (Wasylyshyn et al., 2011). In addition, our work offers a more complete understanding of incentive-based adjustments of cognitive control, which is often studied in paradigms that require frequent adjustments of attention and caution (Botvinick & Braver, 2015; Parro et al., 2018). In this way, our work adds to the growing interest in cognitive control dynamics and offers a route towards a unified view of the costs of cognitive control adjustments, which arise from the need to gradually adjust control, while facing the tradeoff between rapid adjustments and good task performance.

## Methods

### Participants

Across all four studies, we recruited participants with normal or corrected-to-normal vision from Prolific. All participants were fluent English speakers residing in the United States. Prior to the main experiment, participants received instructions on how to perform the Stroop task. They then performed a brief multiple question quiz probing their understanding of the instructions. This quiz served as an attention check and was used to filter out participants who were not attending to the task instructions. Research protocols for all four studies were approved by Brown University’s Institutional Review Board. All of the studies took approximately 1h to complete, and participants were compensated with a fixed rate of $8 per hour. The tasks never included any performance-based monetary incentives. In Study 1 we recruited 48 participants, 4 of which were excluded due to failed attention checks, yielding the final sample of 44 participants (27 female, 14 male, 3 gender not reported; median age = 31). Study 2 included 45 participants, 7 of which were excluded due to failed attention checks, and the final sample consisted of 38 participants (21 female, 17 male; median age = 36). In Study 3 we had 52 participants, 2 of which failed the attention checks, yielding the final sample of 50 participants (17 female, 33 male; median age = 33). Study 4 had 62 participants, 7 of which failed the attention check. The final sample included 55 subjects (17 female, 37 male, 1 gender not reported; median age = 31).

### Task Design

#### Study 1

Participants performed a Stroop task in which they identified the ink color of a color word (e.g., “BLUE”) by pressing one of the 4 keys corresponding to one of the 4 colors (blue, red, yellow, or green). Half of the Stroop trials were either congruent (ink color and the color word are the same) and the other half were incongruent (ink color and the color word are different). Target and distractor colors were randomized under the constraint that both the color word and the ink color did not repeat in two consecutive trials. Participants performed an interval-based version of the Stroop task (Figure 2A) in which they completed as many trials as they wished during a fixed time interval (8-12s). Previous work has used this task to demonstrate reliable adjustments in both drift rates and thresholds in response to incentives, and participant’s performance was well captured by a reward-rate model (Leng et al., 2021). Prior to each interval, participants saw a cue (1.5 s) instructing them to perform the task either as quickly (speed goal) or as accurately (accuracy goal) as possible (Forstmann et al., 2008; Ratcliff & Rouder, 1998). Participants received feedback (1.5 s) on the number of correct responses they made in the current interval. Crucially, there were two types of blocks (Figure 2A) which yielded different levels of control adjustment costs. For half of the blocks, the instructed condition was fixed over the entire block (fixed blocks), and in these blocks participants always received a cue telling them to focus on the same dimension of performance (e.g., speed). The other half of the blocks were varying blocks, in which performance goals changed within a block randomly. Before each block, participants were informed whether this will be a fixed or a varying block, these blocks were intermixed, and their order was counterbalanced across participants. Participants performed 4 blocks, each of which included 20 intervals. Prior to the main task, participants performed practice blocks in which they first practiced the color-button mapping for the Stroop task, and then completed the task with the speed or accuracy condition and got familiarized with the cues for these conditions. We implemented the experiment in Psiturk (Gureckis et al., 2016) and participants performed the task on their own computers and were required to have a keyboard.

#### Study 2

The task was the same as in Study 1, but participants had 3 performance goals: speed, accuracy, and speed/accuracy. In the additional speed/accuracy goal, they were instructed to perform the task as fast and as accurately as possible. The experiment included 6 blocks, each of which included 20 intervals. Half of the blocks were fixed, meaning that participants performed under only one of the 3 goals in each of those blocks (1 block per goal). The other half of the blocks were varying. In these blocks participants switched between the three performance goals randomly.

#### Study 3

Participants performed the same Stroop task described in Study 1. They performed the task as fast (speed goal) or as accurately as possible (accuracy goal), and the task included varying and fixed blocks. In the varying blocks, performance goals switched fully predictably after every interval. Participants completed 6 blocks of this task, and each block consisted of 18 intervals. For both fixed and varying blocks, half of the blocks had a shorter time between the goal cue and task onset, and half of the blocks had longer time. Following the cue presentation (750ms), participants were presented with a fixation cross for either 250ms (short stimulus-onset asynchrony; SOA) or 1000ms (long SOA), immediately followed by task onset.

#### Study 4

Participants performed the same Stroop task as in Study 1, in 5 blocks consisting of 16 intervals. The task included 4 different block types. Participants performed 2 fixed goal blocks (1 block of each goal) in which the performance goal never changed. In varying blocks performance goals changed every 8, 4, or 2 intervals, and participants performed 1 block of each type. Prior to the start of each varying block, participants were informed about the goal switch frequency within the block. Following the cue presentation (1500 ms), participants were presented with a fixation cross for either 1250ms, immediately followed by task onset.

### Statistical Analyses and Drift Diffusion Modeling

#### Behavioral Performance

##### Study 1

To predict reaction times and accuracies on each trial we fitted Bayesian multilevel regressions with goal type (speed vs. accuracy), block type (fixed vs. varying), and their interaction as predictors, while also controlling for the effects of congruency, interval length (8-12s to respond to as many trials as the participant wishes), and the time spent in the interval (time spent within a given interval until the current trial). For the reaction times analyses, we included only correct responses, and we did not analyze responses faster than 250ms and longer than 3000ms. All of the fixed predictors were allowed to vary across subjects. Regression models were fitted in R using the *brms* (Bürkner, 2017; Nalborczyk et al., 2019) package which relies on *Stan* (Carpenter et al., 2017) to implement Markov Chain Monte Carlo (MCMC) algorithm and estimate posterior distributions of parameters. We used ex-Gaussian likelihood to predict reaction times, and Bernoulli to predict accuracy, and the models included uniform priors. For each of the regression models we ran four MCMC simulations (“chains”; 20000 iterations per chain; 19000 warmup), and confirmed convergence by examining trace plots, autocorrelation, variance between and within chains (Gelman & Rubin, 1992), and posterior predictive checks. We summarized the posteriors for each parameter by reporting the mean of the posterior distribution (*b*) and the credible intervals (95% CrI). For inference we relied on the proportion of the posterior samples on one side of 0 (e.g., p*_b_* _< 0_ = 0.01) which represents the probability that given parameter estimate is below 0.

##### Study 2

Models and the model-fitting procedure was the same as in Study 1. The only difference was that the goal type now included 3 levels: speed, accuracy, and speed/accuracy.

##### Study 3

Models and the model-fitting procedure was the same as in Study 1. In addition to the predictors used in Study 1, here we also included the effect of SOA (250ms vs. 1000ms) as the main effect, as well as the interaction between goal type, block type, and SOA.

##### Study 4

Models and the model-fitting procedure was the same as in Study 1. Since this study included 4 different block types (fixed, switch every 8 intervals, switch every 4 intervals, switch every 2 intervals), the effect of block type was coded with an orthogonal polynomial contrast of degree 4. This allowed us to investigate the linear effect of the increase in switch frequency.

#### Drift Diffusion Model

##### Study 1

We fitted hierarchical Bayesian drift diffusion models (Wiecki et al., 2013) to reaction time and accuracy data. For the error trials, we only included trials on which the incorrect response corresponded to the color word and thus driven by the automatic prepotent response to the word. This was done in order to exclude errors which are likely driven by random responding (cf. Leng et al., 2021; Grahek et al., 2023). Fitted models included the effect of congruency on drift rate, as well as the effects of instruction type (speed vs. accuracy), block type (varying vs. fixed), and the interaction between instruction and block type on drift rate. We included the same effects on the threshold, but without the effect of congruency. We also included the effect of running time within interval on the threshold. Finally, we also included group and subject-level parameters of non-decision time, drift rate and non-decision time variability, and the probability of the response coming from another distribution (random response). To ensure the reliability of our findings, we also fitted two additional models: one in which we allowed for non-decision times to vary across different performance goals and block types, and one in which we allowed the performance goal to influence the collapsing bound (Fengler et al., 2021). This analysis showed that the effects we report on drift rate and threshold do not qualitatively change in these more complex models (see Table S2). We ran 5 MCMC simulations (chains; 14000 iterations per chain; 12000 warmup) to estimate the model parameters. We confirmed chain convergence by examining trace plots, the ratio of variances between and within chains (Gelman-Rubin statistic), and posterior predictive checks (Figure S2). As for the regression models, we report the mean of the posterior distribution, the credible intervals, and the probability that the parameter of interest is below 0. When relevant, we compared models using the Deviance Information Criterion (DIC; Spiegelhalter et al., 2002). Based on the Drift Diffusion model estimates, we calculated Euclidian distances between control states under the speed and the accuracy goal. We first range-normalized the posterior estimates by the minimal and maximal values across all goals and blocks. We then calculated the Euclidian distances between the speed and accuracy goals separately in fixed and varying blocks. Subtracting this distance in varying blocks from the distance in fixed blocks provided us with a single metric of control adjustment costs. Larger change in distance indicated higher control adjustment costs. Raw data and analysis scripts are available on GitHub (https://github.com/igrahek/ControlAdjustmentCosts.git). This study was not preregistered.

##### Study 2

The models we used the model-fitting procedure was the same as in Study 1. Since there were 3 different goals in this study, the goal type predictor now included 3 levels: speed, accuracy, and speed/accuracy.

##### Study 3

We used the same models as in Study 1 and fitted them in the same way. We added the additional effect of SOA (250ms vs. 1000ms) as a main effect, as well as the interaction between goal type, block type, and SOA. This was done for both the drift rates and thresholds.

##### Study 4

We used the same model as in Study 1. Since this study included 4 different block types (fixed, switch every 8 intervals, switch every 4 intervals, switch every 2 intervals), the effect of block type was included as a linear, quadratic, and a cubic effect.

### Computational Model

We used the computational model described in the “Control Adjustment Costs: A Dynamical Systems Model” section to simulate performance of an artificial agent in each of the 4 studies. For each of the studies we had our agent perform the task which was closely matched to the task performed by the participants. The agent performed the same number of blocks, block types, performance goals within a block, and the number of intervals in the block as our participants. We report in the results section the average drift rates and thresholds that the model generated for each of the experimental conditions. To simulate task performance, the agent first found the optimal control state (see below) and then gradually moved toward that state (Eq. 1).

#### Determining Optimal Control Intensity

We rely on a reward-rate optimization model (Bogacz et al., 2006; Leng et al., 2021) to specify how optimal control states for each of the performance goals across our experiments are determined. This model (Eq. 3) calculates the expected reward rate (*RR*) given the current value of giving a correct response (*w*_1_) and making an error (*w*_2_). These weights are specified by the current performance goals (e.g., valuing speed over accuracy) or incentives in the environment (e.g., rewards for correct responses). The weights scale the rate of correct responses (1 − *ER*) and the error rate (*ER*) obtained from the Drift Diffusion model (Bogacz et al., 2006). The expected reward rate includes two types of costs. The opportunity cost (Kurzban et al., 2013; Otto & Daw, 2019) which discounts the overall outcome and includes both the decision time (*DT*) and non-decision time (*t*). The second cost is the intensity cost (Shenhav et al., 2013) which represents the quadratic cost on cognitive control intensity, operationalized as the drift rate (*v*). The expected reward rate is thus calculated as:

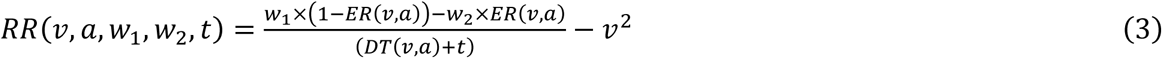

The model finds optimal levels of drift rate (*v*^∗^) and threshold (*a*^∗^) which maximize the expected reward rate:

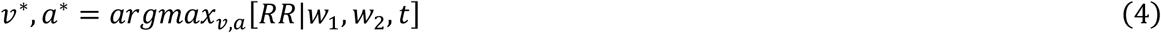

To match our simulations to the range of participant’s performance, we determined the weights for the value of giving a correct response (*w*_1_) and the value of committing an error (*w*_2_) based on the empirical estimates of drift rates, thresholds, and non-decision times in fixed blocks. We employed an inverse reward-rate optimization procedure (cf. Leng et al., 2021) which relies on the reward-rate optimal model (Eq. 3) to find the weights which produce the empirically observed control signals. In this way we determined the two weights for each of the performance goals in each of the studies. On each interval, optimal levels of drift rate (*v*^∗^) and threshold (*a*^∗^) were found by maximizing the expected reward rate given the weights for the current performance goal and the non-decision time (Eq. 3 & 4).

#### Control Adjustment Dynamics

On each interval, after the presentation of the performance cue which determined which weights should be used (e.g., speed vs. accuracy), drift rates and thresholds were gradually adjusted from their starting levels (*v*_0_, *a*_0_), toward the reward-optimal levels (*v*^∗^, *a*^∗^) described above. The initial values of drift rate and threshold (*v* = 3; *a* = 0.8), the speed of adjustment (τ = 0.2; *dt* = 0.005), as well as the noise level (*c* = 0.1) were fixed across all simulations. These parameters were chosen so that the initial values fall within the empirical range of control states and that the control dynamics qualitatively match the observed empirical effects. Due to the use of the sigmoid activation function, drift rates and thresholds were normalized to the 0-1 range before they were adjusted, and they were re-normalized to the original range after the adjustments. The adjustments were performed in discrete steps, and the number of steps was determined by the SOA between cue offset and task onset in each study. At each step, the reward-optimal levels of drift rate and threshold (*v*^∗^, *a*^∗^) were passed through a sigmoid activation function with a fixed gain (*g* = 5) and bias (θ = 0.5).

#### Parameters for Each Study

##### Study 1

We simulated behavior on intervals with the speed goal (*w*_1_ = 8.62, *w*_2_ = 30.39) and the accuracy goal (*w*_1_ = 12.17, *w*_2_ = 916.71). The non-decision time was set to the estimate in the fixed blocks obtained from the drift diffusion model fit (*t* = 0.32). The number of steps for control adjustment was 1500, matched to the SOA in Study 1 (1500 ms). We simulated 4 blocks which included 30 intervals with 10 trials in each interval. Half of the blocks were fixed, meaning that the performance goal, and the corresponding weights, never changed. The other half of blocks included performance goal changes randomly between intervals.

##### Study 2

Behavior was simulated on intervals with the speed goal (*w*_1_ = 11.90, *w*_2_ = 32.98), the accuracy goal (*w*_1_ = 17.88, *w*_2_ = 3673.4), and the speed & goal (*w*_1_ = 15.19, *w*_2_ = 435.97). The non-decision time was set to (*t* = 0.33). The number of steps (1500) was matched to the SOA in Study 2 (1500 ms). We simulated 6 blocks which included 20 intervals with 10 trials in each interval. Half of the blocks were fixed, meaning that the performance goal, and the corresponding weights, never changed. The other half of blocks included performance goal changes randomly between intervals.

##### Study 3

Simulations included the speed goal (*w*_1_ = 15.65, *w*_2_ = 145.76) and the accuracy goal (*w*_1_ = 16.58, *w*_2_ = 671.97). The non-decision time was set to *t* = 0.39 . The number of steps was matched to the time between the cue onset and the first trial onset in Study 3 (short SOA = 1000 ms; long SOA = 1750 ms). We simulated 6 blocks which included 18 intervals with 10 trials in each interval. Half of the blocks were fixed, meaning that the performance goal, and the corresponding weights, never changed. The other half of blocks included changes between the two performance goals every 1 interval.

##### Study 4

This simulation included two performance goals: the speed goal (*w*_1_ = 17.79, *w*_2_ = 136.07) and the accuracy goal (*w*_1_ = 20.50, *w*_2_ = 1787.4). The non-decision time was set to: *t* = 0.38. The number of steps for control adjustment was 2750 matched the time from cue onset until the first trial onset in Study 4 (2750 ms). We simulated 5 blocks which included 16 intervals with 10 trials in each interval. The blocks were divided into 4 groups: fixed blocks (2 blocks; always the same performance goal), switch every 8 intervals (1 block), switch every 4 intervals (1 block), and switch every 2 intervals (1 block). The gain parameters differed across the fixed blocks (*g* = 5), and the switch 8 (*g* = 3), switch 4 (*g* = 2.5), and switch 2 blocks (*g* = 2).

## Supplementary materials

**Table S1.** Study 1 parameter estimates for hierarchical regressions predicting behavioral performance (reaction times and accuracy) and parameter estimates of the Drift Diffusion Model (a – threshold; v – drift rate; t – non-decision time; z – bias).

| *Parameter* | *Estimate* | *95% Credible Interval* | *Posterior probability (p<0)* |
| --- | --- | --- | --- |
| *Reaction times (ms)* |  |  |  |
| Intercept | 718.62 | 682.05, 755.69 | 0.00 |
| Congruency (I-C) | 39.18 | 28.82, 49.61 | 0.00 |
| Goal (A-S) | 128.25 | 85.36, 172.52 | 0.00 |
| Block (V-F) | -12.09 | -27.66, 4.04 | 0.93 |
| Goal * Block | 28.09 | 3.54, 51.41 | 0.01 |
| Interval number | -7.72 | -11.48, -3.92 | 1.00 |
| Interval length | 3.81 | 2.01, 5.58 | 0.00 |
| Time in interval | -22.71 | -24.68, -20.74 | 1.00 |
| *Accuracy (log odds)* |  |  |  |
| Intercept | 1.66 | 1.35, 1.99 | 0.00 |
| Congruency (I-C) | -0.25 | -0.36, -0.14 | 1.00 |
| Goal (A-S) | 1.67 | 1.12, 2.22 | 0.00 |
| Block (V-F) | -0.11 | -0.29, 0.07 | 0.89 |
| Goal * Block | 0.44 | 0.12, 0.76 | 0.00 |
| Interval number | -0.17 | -0.23, -0.12 | 1.00 |
| Interval length | -0.03 | -0.05, 0.00 | 0.96 |
| Time in interval | -0.01 | -0.04, 0.02 | 0.80 |
| *Drift Diffusion Model Estimates* |  |  |  |
| (a) Intercept | 0.94 | 0.87, 1.01 | 0.00 |
| (a) Goal (A-S) | 0.42 | 0.27, 0.58 | 0.00 |
| (a) Block (V-F) | 0.00 | -0.08, 0.08 | 0.49 |
| (a) Goal * Block | 0.08 | 0.03, 0.13 | 0.00 |
| (a) Time in interval | -0.07 | -0.08, -0.06 | 0.00 |
| (v) Intercept | 2.66 | 2.49, 2.84 | 0.00 |
| (v) Goal (A-S) | 0.54 | 0.31, 0.77 | 0.00 |
| (v) Block (V-F) | 0.03 | -0.16, 0.09 | 0.29 |
| (v) Goal * Block | 0.05 | -0.12, 0.21 | 0.28 |
| (v) Congruency (I-C) | -0.34 | -0.41, -0.28 | 0.00 |
| (t) Intercept | 0.32 | 0.3, 0.34 | 0.00 |
| (z) Congruency (I-C) | -0.02 | -0.03, -0.01 | 0.00 |
| Lapse rate | 0.00 | 0.00, 0.00 | 0.00 |

**Table S2.** Study 1 parameter estimates for parameter estimates of the Drift Diffusion Models which include:1) effects on non-decision time, and 2) a collapsing bound (a – threshold; v – drift rate; t – non-decision time; z – bias).

| *Parameter* | *Estimate* | *95% Credible Interval* | *Posterior probability ($p < 0$)* |
| --- | --- | --- | --- |
| *Drift Diffusion Model Estimates with Effects on non-decision time* |  |  |  |
| (a) Intercept | 0.86 | 0.80, 0.93 | 0.00 |
| (a) Goal (S-A) | 0.30 | 0.18, 0.41 | 0.00 |
| (a) Block (V-F) | 0.00 | -0.06, 0.06 | 0.44 |
| (a) Goal * Block | 0.05 | 0.01, 0.10 | 0.01 |
| (a) Time in interval | -0.07 | -0.08, -0.06 | 0.00 |
| (v) Intercept | 2.38 | 2.20, 2.56 | 0.00 |
| (v) Goal (A-S) | 0.51 | 0.30, 0.73 | 0.00 |
| (v) Block (V-F) | -0.03 | -0.14, 0.10 | 0.34 |
| (v) Goal * Block | 0.04 | -0.12, 0.20 | 0.29 |
| (v) Congruency (I-C) | -0.31 | -0.37, -0.25 | 0.00 |
| (t) Intercept | 0.34 | 0.32, 0.37 | 0.00 |
| (t) Goal (A-S) | 0.04 | 0.02, 0.07 | 0.00 |
| (t) Block (V-F) | 0.00 | -0.02, 0.02 | 0.41 |
| (t) Goal * Block | 0.01 | 0.00, 0.03 | 0.10 |
| (z) Congruency (I-C) | -0.03 | -0.04, 0.02 | 0.00 |
| Lapse rate | 0.00 | 0.00, 0.00 | 0.00 |
| *Drift Diffusion Model Estimates with a Collapsing Bound* |  |  |  |
| (a) Intercept | 0.55 | 0.50, 0.60 | 0.00 |
| (a) Goal (A-S) | 0.20 | 0.12, 0.28 | 0.00 |
| (a) Block (V-F) | 0.00 | -0.04, 0.05 | 0.46 |
| (a) Goal * Block | 0.04 | 0.01, 0.07 | 0.00 |
| (v) Intercept | 1.87 | 1.72, 2.02 | 0.00 |
| (v) Goal (A-S) | 0.50 | 0.31, 0.68 | 0.00 |
| (v) Block (V-F) | -0.02 | -0.11, 0.08 | 0.36 |
| (v) Goal * Block | 0.05 | -0.07, 0.17 | 0.20 |
| (v) Congruency (I-C) | -0.34 | -0.39, -0.30 | 0.00 |
| (t) Intercept | 0.26 | 0.23, 0.28 | 0.00 |
| (t) Goal (S-A) | -0.02 | -0.03, -0.01 | 0.00 |
| (theta) Intercept | 0.16 | -0.08, 0.34 | 0.12 |
| (theta) Goal (S-A) | -0.10 | -0.19, -0.01 | 0.01 |
| Lapse rate | 0.01 | 0.01, 0.01 | 0.00 |

**Table S3.**
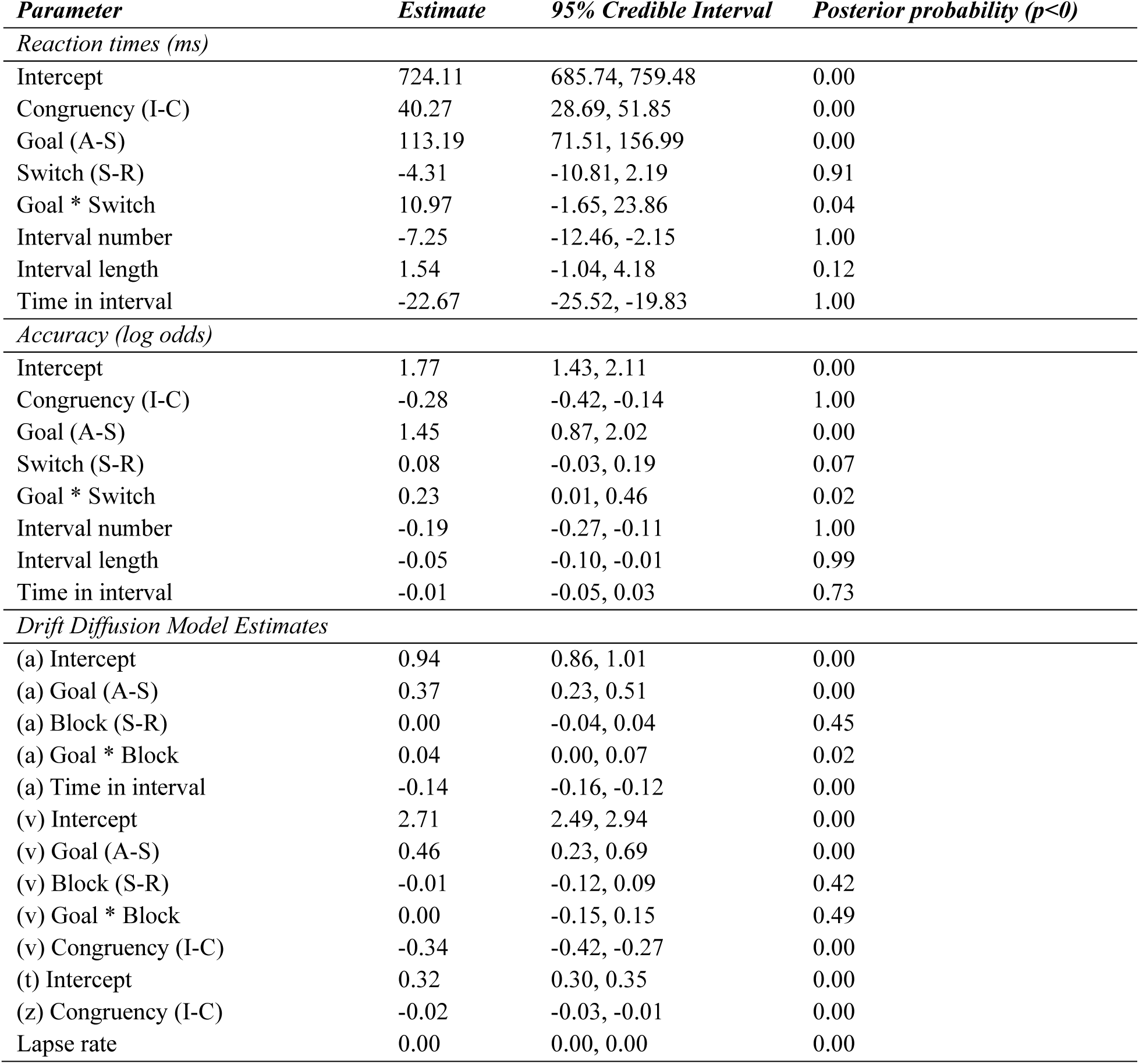
Study 1 – switch vs. repeat intervals analysis. Parameter estimates for hierarchical regressions predicting behavioral performance (reaction times and accuracy) and parameter estimates of the Drift Diffusion Model (a – threshold; v – drift rate; t – non-decision time; z – bias).

**Table S4.** Study 1 parameter estimates for hierarchical regressions predicting behavioral performance (reaction times and accuracy) with considering control adjustment costs across congruency levels

| *Parameter* | *Estimate* | *95% Credible Interval* | *Posterior probability (p<0)* |
| --- | --- | --- | --- |
| *Reaction times (ms)* |  |  |  |
| Intercept | 717.45 | 684.19, 749.77 | 0.00 |
| Congruency (I-C) | 35.17 | 25.33, 45.10 | 0.00 |
| Goal (A-S) | 126.22 | 84.39, 168.50 | 0.00 |
| Block (V-F) | -7.29 | -21.84, 7.77 | 0.83 |
| Congruency * Goal | 21.14 | 9.42, 33.34 | 0.00 |
| Congruency * Block | -2.69 | -12.63, 6.80 | 0.70 |
| Goal * Block | 27.49 | 4.74, 50.55 | 0.01 |
| Congruency * Goal * Block | 21.16 | 3.14, 39.25 | 0.01 |
| Interval number | -6.51 | -10.25, -2.76 | 1.00 |
| Interval length | 3.77 | 1.91, 5.55 | 0.00 |
| Time in interval | -22.80 | -24.80, -20.73 | 1.00 |
| *Accuracy (log odds)* |  |  |  |
| Intercept | 1.67 | 1.36, 2.00 | 0.00 |
| Congruency (I-C) | -0.27 | -0.38, -0.16 | 1.00 |
| Goal (A-S) | 1.70 | 1.14, 2.24 | 0.00 |
| Block (V-F) | 0.00 | -0.17, 0.17 | 0.50 |
| Congruency * Goal | -0.04 | -0.20, 0.11 | 0.71 |
| Congruency * Block | 0.01 | -0.14, 0.15 | 0.45 |
| Goal * Block | 0.46 | 0.15, 0.77 | 0.00 |
| Congruency * Goal * Block | -0.23 | -0.51, 0.05 | 0.96 |
| Interval number | -0.13 | -0.18, -0.07 | 1.00 |
| Interval length | -0.03 | -0.05, 0.00 | 0.96 |
| Time in interval | -0.01 | -0.04, 0.02 | 0.78 |

**Table S5.** Study 2 parameter estimates for hierarchical regressions predicting behavioral performance (reaction times and accuracy) and parameter estimates of the Drift Diffusion Model (a – threshold; v – drift rate; t – non-decision time; z – bias).

| *Parameter* | *Estimate* | *95% Credible Interval* | *Posterior probability ($p < 0$)* |
| --- | --- | --- | --- |
| *Reaction times (ms)* |  |  |  |
| Intercept | 669.25 | 641.01, 697.21 | 0.00 |
| Congruency (I-C) | 39.30 | 31.53, 47.01 | 0.00 |
| Goal (A-S) | 143.32 | 95.18, 190.48 | 0.00 |
| Goal (A-S/A) | -52.74 | -79.10, -25.10 | 1.00 |
| Block (V-F) | -3.17 | -17.84, 11.35 | 0.67 |
| Goal (A-S) * Block | 30.30 | -3.83, 64.80 | 0.04 |
| Goal (A-S/A) * Block | -8.43 | -35.40, 17.49 | 0.74 |
| Interval number | -9.20 | -12.91, -5.50 | 1.00 |
| Interval length | 1.91 | 0.18, 3.65 | 0.02 |
| Time in interval | -21.74 | -23.71, -19.81 | 1.00 |
| *Accuracy (log odds)* |  |  |  |
| Intercept | 1.83 | 1.45, 2.21 | 0.00 |
| Congruency (I-C) | -0.32 | -0.50, -0.14 | 1.00 |
| Goal (S-A) | 2.01 | 1.50, 2.52 | 0.00 |
| Goal (A-S/A) | -0.69 | -0.94, -0.45 | 1.00 |
| Block (V-F) | -0.06 | -0.23, 0.12 | 0.75 |
| Goal (A-S) * Block | 0.75 | 0.31, 1.20 | 0.00 |
| Goal (A-S/A) * Block | -0.27 | -0.64, 0.10 | 0.92 |
| Interval number | 0.01 | -0.04, 0.07 | 0.33 |
| Interval length | -0.02 | -0.05, 0.01 | 0.92 |
| Time in interval | -0.06 | -0.09, -0.03 | 1.00 |
| *Drift Diffusion Model Estimates* |  |  |  |
| (a) Intercept | 0.89 | 0.82, 0.97 | 0.00 |
| (a) Goal (A-S) | 0.48 | 0.38, 0.58 | 0.00 |
| (a) Goal (A-S/A) | 0.20 | 0.10, 0.31 | 0.00 |
| (a) Block (V-F) | 0.00 | -0.06, 0.06 | 0.51 |
| (a) Goal (A-S) * Block | 0.11 | 0.04, 0.18 | 0.00 |
| (a) Goal (A-S/A) * Block | 0.04 | -0.03, 0.11 | 0.17 |
| (a) Time in interval | -0.07 | -0.08, -0.05 | 0.00 |
| (v) Intercept | 3.20 | 2.91, 3.49 | 0.00 |
| (v) Goal (A-S) | 0.66 | 0.47, 0.86 | 0.00 |
| (v) Goal (A-S/A) | 0.27 | 0.09, 0.46 | 0.01 |
| (v) Block (V-F) | 0.07 | -0.06, 0.20 | 0.18 |
| (v) Goal (A-S) * Block | 0.36 | 0.16, 0.55 | 0.00 |
| (v) Goal (A-S/A) * Block | 0.09 | -0.09, 0.27 | 0.21 |
| (v) Congruency (I-C) | -0.41 | -0.50, -0.33 | 0.00 |
| (t) Intercept | 0.33 | 0.31, 0.36 | 0.00 |
| (z) Congruency (I-C) | -0.02 | -0.03, -0.02 | 0.00 |
| Lapse rate | 0.00 | 0.00, 0.00 | 0.00 |

**Figure S1.**
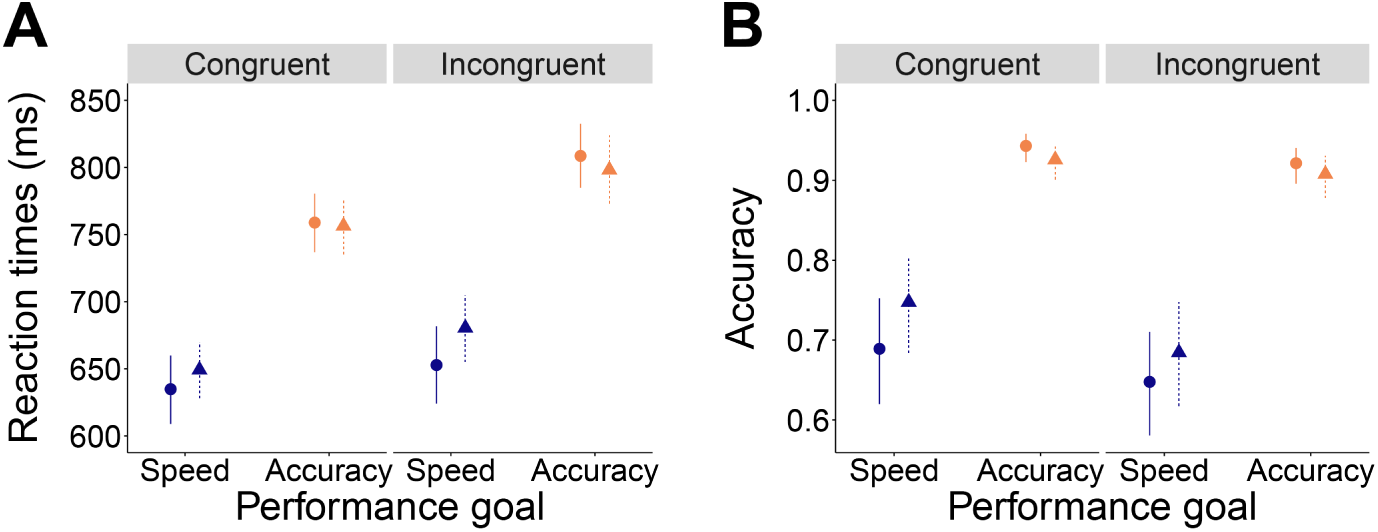
Control adjustment costs in congruent and incongruent trials in Study 1. **A.** While control adjustment costs are present in both congruent and incongruent trials when analyzing reaction times, they are more pronounced in incongruent trials. **B.** When analyzing accuracy, control adjustment costs are present in both congruent and incongruent trials, but are more pronounced in congruent trials.

**Table S6.**
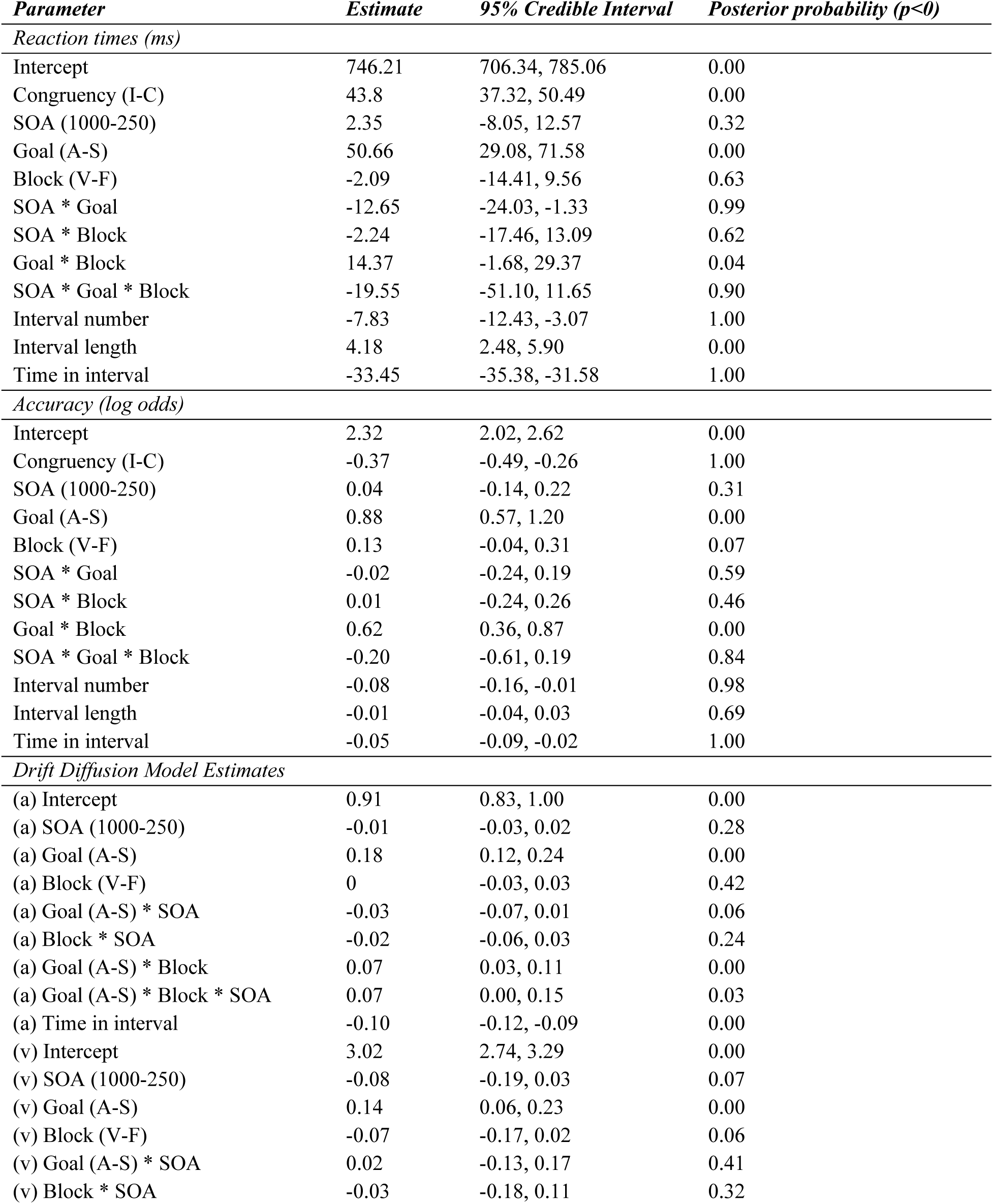
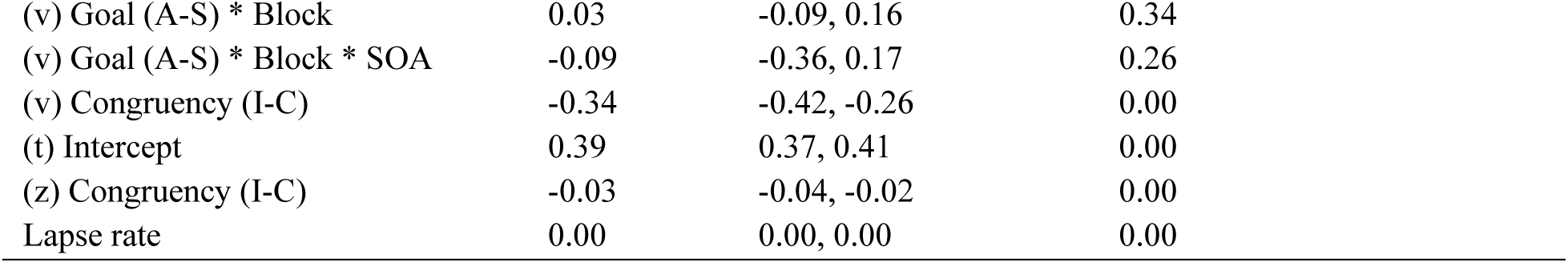
Study 3 parameter estimates for hierarchical regressions predicting behavioral performance (reaction times and accuracy) and parameter estimates of the Drift Diffusion Model (a – threshold; v – drift rate; t – non-decision time; z – bias).

| *Parameter* | *Estimate* | *95% Credible Interval* | *Posterior probability ($p < 0$)* |
| --- | --- | --- | --- |
| *Reaction times (ms)* |  |  |  |
| Intercept | 746.21 | 706.34, 785.06 | 0.00 |
| Congruency (I-C) | 43.8 | 37.32, 50.49 | 0.00 |
| SOA (1000-250) | 2.35 | -8.05, 12.57 | 0.32 |
| Goal (A-S) | 50.66 | 29.08, 71.58 | 0.00 |
| Block (V-F) | -2.09 | -14.41, 9.56 | 0.63 |
| SOA * Goal | -12.65 | -24.03, -1.33 | 0.99 |
| SOA * Block | -2.24 | -17.46, 13.09 | 0.62 |
| Goal * Block | 14.37 | -1.68, 29.37 | 0.04 |
| SOA * Goal * Block | -19.55 | -51.10, 11.65 | 0.90 |
| Interval number | -7.83 | -12.43, -3.07 | 1.00 |
| Interval length | 4.18 | 2.48, 5.90 | 0.00 |
| Time in interval | -33.45 | -35.38, -31.58 | 1.00 |
| *Accuracy (log odds)* |  |  |  |
| Intercept | 2.32 | 2.02, 2.62 | 0.00 |
| Congruency (I-C) | -0.37 | -0.49, -0.26 | 1.00 |
| SOA (1000-250) | 0.04 | -0.14, 0.22 | 0.31 |
| Goal (A-S) | 0.88 | 0.57, 1.20 | 0.00 |
| Block (V-F) | 0.13 | -0.04, 0.31 | 0.07 |
| SOA * Goal | -0.02 | -0.24, 0.19 | 0.59 |
| SOA * Block | 0.01 | -0.24, 0.26 | 0.46 |
| Goal * Block | 0.62 | 0.36, 0.87 | 0.00 |
| SOA * Goal * Block | -0.20 | -0.61, 0.19 | 0.84 |
| Interval number | -0.08 | -0.16, -0.01 | 0.98 |
| Interval length | -0.01 | -0.04, 0.03 | 0.69 |
| Time in interval | -0.05 | -0.09, -0.02 | 1.00 |
| *Drift Diffusion Model Estimates* |  |  |  |
| (a) Intercept | 0.91 | 0.83, 1.00 | 0.00 |
| (a) SOA (1000-250) | -0.01 | -0.03, 0.02 | 0.28 |
| (a) Goal (A-S) | 0.18 | 0.12, 0.24 | 0.00 |
| (a) Block (V-F) | 0 | -0.03, 0.03 | 0.42 |
| (a) Goal (A-S) * SOA | -0.03 | -0.07, 0.01 | 0.06 |
| (a) Block * SOA | -0.02 | -0.06, 0.03 | 0.24 |
| (a) Goal (A-S) * Block | 0.07 | 0.03, 0.11 | 0.00 |
| (a) Goal (A-S) * Block * SOA | 0.07 | 0.00, 0.15 | 0.03 |
| (a) Time in interval | -0.10 | -0.12, -0.09 | 0.00 |
| (v) Intercept | 3.02 | 2.74, 3.29 | 0.00 |
| (v) SOA (1000-250) | -0.08 | -0.19, 0.03 | 0.07 |
| (v) Goal (A-S) | 0.14 | 0.06, 0.23 | 0.00 |
| (v) Block (V-F) | -0.07 | -0.17, 0.02 | 0.06 |
| (v) Goal (A-S) * SOA | 0.02 | -0.13, 0.17 | 0.41 |
| (v) Block * SOA | -0.03 | -0.18, 0.11 | 0.32 |
| (v) Goal (A-S) * Block | 0.03 | -0.09, 0.16 | 0.34 |
| (v) Goal (A-S) * Block * SOA | -0.09 | -0.36, 0.17 | 0.26 |
| (v) Congruency (I-C) | -0.34 | -0.42, -0.26 | 0.00 |
| (t) Intercept | 0.39 | 0.37, 0.41 | 0.00 |
| (z) Congruency (I-C) | -0.03 | -0.04, -0.02 | 0.00 |
| Lapse rate | 0.00 | 0.00, 0.00 | 0.00 |

**Table S7.** Study 4 parameter estimates for hierarchical regressions predicting behavioral performance (reaction times and accuracy) and parameter estimates of the Drift Diffusion Model (a – threshold; v – drift rate; t – non-decision time; z – bias).

| *Parameter* | *Estimate* | *95% Credible Interval* | *Posterior probability (p<0)* |
| --- | --- | --- | --- |
| *Reaction times (ms)* |  |  |  |
| Intercept | 728.8 | 683.41, 772.02 | 0.00 |
| Congruency (I-C) | 40.04 | 32.34, 47.86 | 0.00 |
| Goal (A-S) | 54.14 | 37.96, 70.30 | 0.00 |
| Switch frequency (linear effect) | -3.01 | -16.15, 9.71 | 0.67 |
| Switch frequency (quadratic effect) | -6.73 | -17.44, 4.39 | 0.90 |
| Switch frequency (cubic effect) | 4.39 | -3.51, 11.95 | 0.12 |
| Switch frequency (linear effect) *Goal | 25.20 | 8.50, 42.58 | 0.00 |
| Switch frequency (quadratic effect) *Goal | 20.98 | 8.22, 33.39 | 0.00 |
| Switch frequency (cubic effect) *Goal | 3.03 | -8.45, 14.90 | 0.31 |
| Interval number | -12.89 | -17.33, -8.62 | 1.00 |
| Interval length | 2.94 | 1.72, 4.18 | 0.00 |
| Time in interval | -13.76 | -15.03, -12.47 | 1.00 |
| *Accuracy (log odds)* |  |  |  |
| Intercept | 2.49 | 2.25, 2.70 | 0.00 |
| Congruency (I-C) | -0.25 | -0.35, -0.15 | 1.00 |
| Goal (A-S) | 0.80 | 0.58, 1.04 | 0.00 |
| Switch frequency (linear effect) | -0.24 | -0.39, -0.10 | 1.00 |
| Switch frequency (quadratic effect) | -0.20 | -0.35, -0.07 | 1.00 |
| Switch frequency (cubic effect) | 0.01 | -0.06, 0.09 | 0.35 |
| Switch frequency (linear effect) * Goal | 0.06 | 0.00, 0.12 | 0.03 |
| Switch frequency (quadratic effect) * Goal | -0.03 | -0.06, 0.00 | 0.96 |
| Switch frequency (cubic effect) * Goal | -0.09 | -0.12, -0.07 | 1.00 |
| Interval number | 0.48 | 0.25, 0.74 | 0.00 |
| Interval length | 0.21 | 0.00, 0.42 | 0.03 |
| Time in interval | 0.09 | -0.09, 0.27 | 0.15 |
| *Drift Diffusion Model Estimates* |  |  |  |
| (a) Intercept | 0.96 | 0.89, 1.03 | 0.00 |
| (a) Goal (A-S) | 0.23 | 0.17, 0.30 | 0.00 |
| (a) Switch frequency (linear effect) | -0.05 | -0.08, -0.02 | 0.00 |
| (a) Switch frequency (quadratic effect) | -0.06 | -0.09, -0.03 | 0.00 |
| (a) Switch frequency (cubic effect) | 0.00 | -0.03, 0.02 | 0.47 |
| (a) Switch frequency (linear effect) * Goal | -0.12 | -0.21, -0.04 | 0.00 |
| (a) Switch frequency (quadratic effect) * Goal | -0.02 | -0.04, 0.00 | 0.06 |
| (a) Switch frequency (cubic effect) * Goal | 0.00 | -0.02, 0.02 | 0.47 |
| (a) Time in interval | -0.05 | -0.06, -0.04 | 0.00 |
| (v) Intercept | 3.38 | 3.18, 3.57 | 0.00 |
| (v) Goal (A-S) | 0.30 | 0.17, 0.44 | 0.00 |
| (v) Switch frequency (linear effect) | 0.01 | -0.08, 0.10 | 0.41 |
| (v) Switch frequency (quadratic effect) | -0.08 | -0.16, 0.00 | 0.02 |
| (v) Switch frequency (cubic effect) | -0.01 | -0.08, 0.05 | 0.34 |
| (v) Switch frequency (linear effect) * Goal | -0.20 | -0.43, 0.05 | 0.05 |
| (v) Switch frequency (quadratic effect) * Goal | 0.01 | -0.05, 0.07 | 0.39 |
| (v) Switch frequency (cubic effect) * Goal | 0.05 | -0.01, 0.11 | 0.06 |
| (v) Congruency (I-C) | -0.38 | -0.45, -0.32 | 0.00 |
| (t) Intercept | 0.38 | 0.36, 0.41 | 0.00 |
| (z) Congruency (I-C) | -0.02 | -0.03, -0.01 | 0.00 |
| Lapse rate | 0.00 | 0.00, 0.00 | 0.00 |

**Figure S2.**
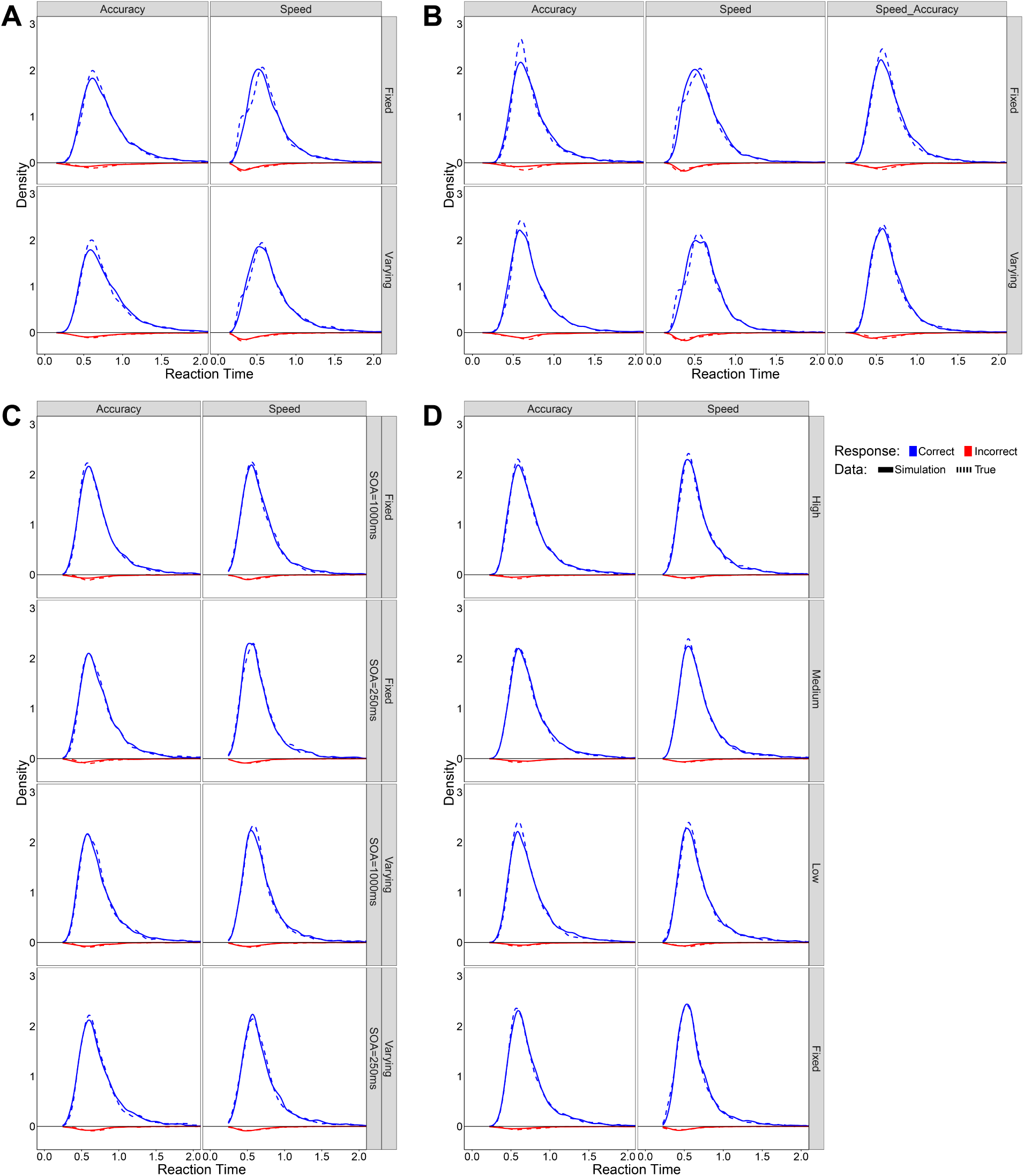
Posterior predictive checks for the fitted Drift Diffusion models. Distributions of simulated reaction times from posterior predictive checks (solid lines) match the empirical reaction time distributions (dashed lines) for both correct and incorrect responses. This is true for Study 1 (**A**), Study 2 (**B**), Study 3 (**C**), and Study 4 (**D**).

## Footnotes

1 All of the estimates for accuracy analyses are in log odds.

